# Rotational Dynamics Reduce Interference Between Sensory and Memory Representations

**DOI:** 10.1101/641159

**Authors:** Alexandra Libby, Timothy J. Buschman

## Abstract

Sensory stimuli arrive in a continuous stream. By learning statistical regularities in the sequence of stimuli, the brain can predict future stimuli (Xu et al. 2012; Gavornik and Bear 2014; Maniscalco et al. 2018; J. Fiser and Aslin 2002). Such learning requires associating immediate sensory information with the memory of recently encountered stimuli (Ostojic and Fusi 2013; Kiyonaga et al. 2017). However, new sensory information can also interfere with short-term memories (Parthasarathy et al. 2017). How the brain prevents such interference is unknown. Here, we show that sensory representations rotate in neural space over time, to form an independent memory representation, thus reducing interference with future sensory inputs. We used an implicit learning paradigm in mice to study how statistical regularities in a sequence of stimuli are learned and represented in primary auditory cortex. Mice experienced both common sequences of stimuli (e.g. ABCD) and uncommon sequences (e.g. XYCD). Over four days of learning, the neural population representation of commonly associated stimuli (e.g. A and C) converged. This facilitated the prediction of upcoming stimuli, but also led unexpected sensory inputs to overwrite the sensory representation of previous stimuli (postdiction). Surprisingly, we found the memory of previous stimuli persisted in a second, orthogonal dimension. Unsupervised clustering of functional cell types revealed that the emergence of this second memory dimension is supported by two separate types of neurons; a ‘stable’ population that maintained its selectivity throughout the sequence and a ‘switching’ population that dynamically inverted its selectivity. This combination of sustained and dynamic representations produces a rotation of the encoding dimension in the neural population. This rotational dynamic may be a general principle, by which the cortex protects memories of prior events from interference by incoming stimuli.

## Main Text

Predictions are a key neural computation; they improve stimulus processing (de Lange, Heilbron, and Kok 2018; Brandman and Peelen 2017; A. Fiser et al. 2016) and speed behavioral responses (Jaramillo and Zador 2011; Chun and Jiang 1998; Dehaene et al. 2015). Predictions are based on experience: we use previously learned cross-temporal associations to anticipate future stimuli (Ostojic and Fusi 2013). These associations can be learned from statistical regularities in a sequence of stimuli, without supervision or an explicit task (Gavornik and Bear 2014; Li and DiCarlo 2008, 2012; Maheu, Dehaene, and Meyniel 2019; Kim et al. 2009). Yet, despite its importance, the neural mechanisms underlying both the learning and representation of predictions is largely unknown. In particular, it is unknown how the brain simultaneously represents both current and previous sensory inputs to allow associations to be learned across time (Kiyonaga et al. 2017).

To address this, we recorded from auditory cortex in mice, while they gained experience with sequences of four chords (Fig. 1A, see Supporting Information, SI). Statistical regularities in the transitions between chords created learnable predictions within the sequences (Fig. 1B). Specifically, sequences began with one of two contextual chord pairs, either an ‘A’ and ‘B’ stimulus pair (the AB context) or an ‘X’ and ‘Y’ stimulus pair (the XY context). These contexts predicted what chord would follow: on 68% of trials, the AB context was followed by a C chord and the XY context was followed by a C* chord. All sequences ended with a D chord. However, on a subset of trials (20%), the animals unexpectedly heard the other stimulus (i.e. rare sequences: ABC*D and XYCD; the remaining 12% of trials were ambiguous stimuli, not analyzed here). The overall likelihood of each sensory stimulus was balanced across conditions. Therefore, at the start of each sequence, the animal had no *a priori* expectation as to what chords it would experience. Only after the presentation of the contextual stimuli (AB/XY) could the animal predict the upcoming C/C* stimulus. Beginning naïve, the animals experienced 1500 sequences of stimuli per day, for 4 consecutive days (no behavioral response was required; see SI).

**Figure 1.**
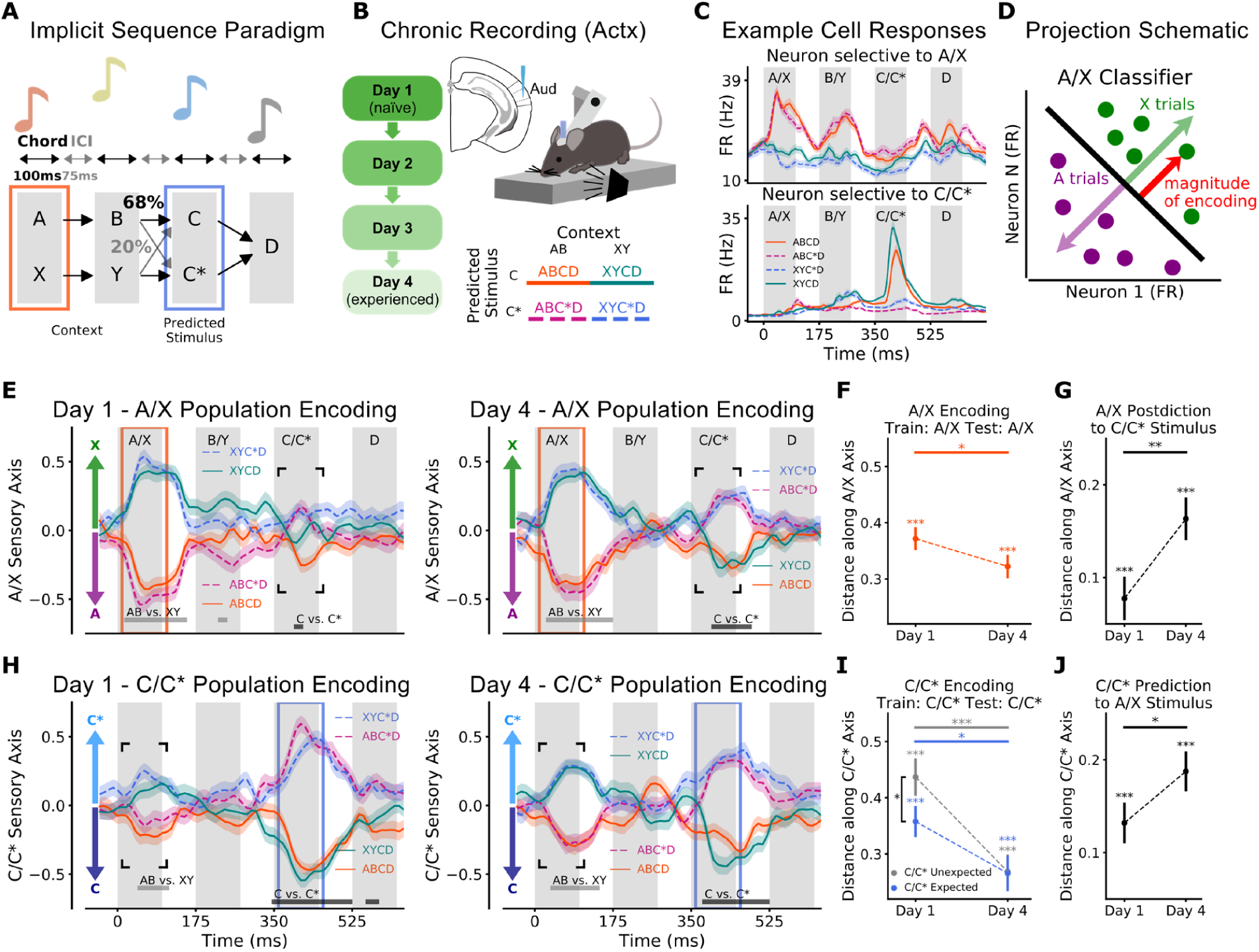
Neural Encoding of Statistical Regularities in Sensory Inputs. **(A)** Schematic of implicit sequence learning paradigm. Animals heard 1500 sequences of auditory chords every day. Statistical regularities were embedded in the sequences: 68% expected trials (ABCD, XYC*D), 20% unexpected (ABC*D, XYCD), 12% mixed stimuli (not analyzed here; see SI for details). **(B)** Silicon probes were chronically implanted in right auditory cortex. Animals experienced sequences over 4 days. All mice were initially naïve to sequences. Legend of four trial types (colors maintained throughout the manuscript). **(C)** Response of two example neurons to auditory sequences preferring AB context (top) or C stimulus (bottom). Gray bars indicate stimulus periods. **(D)** Schematic of N-Dimensional firing rate activity projected onto encoding axis. Axis is defined as normal to trained classifiers (see SI for details). **(E)** Population encoding of A/X information across the trial on Day 1 (left) and Day 4 (right). Firing rate activity was projected onto the A/X sensory axis of each individual animal (training period outlined in orange), z-scored and combined across animals. Mean and s.e.m of projections shown across time for all four trial types. Positive projections and negative projections indicate X (green) and A (purple) encoding, respectively. Light and dark grey bars mark significant differences for AB vs XY and C vs. C*, respectively (p ≤ 0.001, t-test, Bonferroni corrected). **(F)** Encoding of A/X stimulus along the A/X sensory axis (orange box in E). Mean and s.e.m. of absolute distance along axis combined across trials. Absolute distance allows conditions to be combined as negatively encoded conditions are flipped. Thus, positive values indicate correct encoding strength. **(G)** A/X postdiction in response to C/C* stimulus (C/C* period marked by black box in E). For example, C* trials were coded as X on A/X sensory axis. **(H)** Population encoding of C/C* information across the trial on Day 1 and 4 (as in **D**). Blue box marks training period of C/C* sensory axis. Positive projections and negative projections indicate C* (light blue) and C (dark blue) encoding. **(I)** C/C* encoding along the C/C* sensory axis (as in **F**). Trials were divided by expected (ABCD, XYC*D) and unexpected (ABC*D, XYCD). **(J)** C/C* prediction in response to A/X stimulus (A/X period marked by black box in H). For example, X trials were coded as C* on C/C* sensory axis. For all panels, p-values: * ≤ 0.05, ** ≤ 0.01, *** ≤ 0.001

To track how expectation shaped neural responses, we recorded 522 neurons from the auditory cortex of 7 mice, over the 4 days of sequence experience (see SI). Although all recordings were chronic, individual neurons were not tracked across recording sessions. Individual neurons showed selectivity to both context chords (A vs. X and B vs. Y) and the predicted stimulus (C vs. C*; see Fig. 1C). To measure the stimulus representation in the neural population on each day, we defined an encoding axis for each chord in the sequence. The encoding axis was defined as the vector normal to the hyperplane that best classified the population response to each stimulus pair (Fig. 1D, classifier was a linear SVM trained on average activity from 10-110 ms after stimulus onset, see SI for details and Fig. S2 for classifier performance). To ensure an unbiased classifier, trial types were balanced (i.e. an equal number of ABCD, ABC*D, XYCD, and XYC*D trials) and all analyses were performed on withheld data. To measure population encoding of each stimulus, we projected neural activity onto the encoding axis (Fig. 1D). These signed projections represent the relative encoding strength of each stimulus (e.g. X is positive and A is negative along the A/X axis, as seen in Fig. 1E).

Figure 1E shows the temporal evolution of A/X sensory encoding. As expected, the population strongly encoded the A/X stimulus during its presentation (Fig. 1E, orange shaded region; average encoding during presentation, D1=0.37, D4=0.32, p<1/5000, bootstrap test, Fig. 1F). However, this information quickly decayed after the sensory stimulus. The A/X sensory classifier only weakly decoded the B/Y stimulus (Fig. S4), suggesting the encoding of A/X sensory information is independent from B/Y representations.

C/C* stimuli were also strongly represented during their presentation (Fig. 1H, blue shaded region). Similar to previous studies examining responses to rare stimuli (Kurkela et al. 2018; Chen, Helmchen, and Lütcke 2015; Natan et al. 2015), unexpected C/C* stimuli were more strongly encoded than expected stimuli on day 1 (Fig. 1I; D1 difference=0.079 p=0.033, permutation test). Importantly, because our paradigm was balanced, the difference between the expected and unexpected responses could not be attributed to the stimulus, but solely to its contextually-defined expectation within the sequence. Consistent with adaptation (F. P. de Lange, Heilbron, and Kok 2018; Kato, Gillet, and Isaacson 2015), encoding strength of both expected and unexpected trials decreased over days (Fig. 1K; expected D4-D1 = −0.092, p=0.016; unexpected D4-D1 = −0.17, p=0.0002, permutation test).

In addition, there was predictive encoding of the expected C/C* stimulus during the presentation of A/X: when A or X was presented, the neural population began to encode C or C*, respectively (Fig. 1H, black box, and Fig. 1J; D1 = 0.13, p<1/5000; D4 = 0.19, p<1/5000, permutation test). This effect grew with experience (D4-D1 = 0.056, p = 0.038, permutation test). Indeed, by day 4, the predicted response of C/C* during A/X was nearly as strong as the sensory response to C/C* stimuli themselves.

To directly demonstrate the relationship between associated stimuli, we plotted the population activity in a 2D state space defined by the A/X sensory and C/C* sensory encoding axes. These axes are independent (but not necessarily orthogonal as we show below) and so this space allows us to track the evolution of both A/X sensory and C/C* sensory information during the sequence. On day 1, neural activity, during the presentation of the A/X stimulus, evolved along the A/X axis, reflecting the encoding of A/X sensory information in the neural population (Fig. 2A, left). However, by day 4, the neural activity evoked by the A/X stimulus evolved along a rotated axis (Fig. 2A, right). Now, the A/X stimulus induced encoding of their predicted stimuli (C and C* for A and X, respectively). Predictions increased with experience, reflected in the increased angle of the first principle component (PC) of the neural trajectory over days (Fig. 2A, insets, and Fig. 2B, D1=19 deg, D4=31 deg. D4-D1=12 deg, p≤1/5000, permutation test, see SI for details). Predictions were also observed immediately before the onset of the C/C* stimulus on day 4 (Fig. 2D, square points are before onset of C/C*; distance along C/C* axis from 300-350 ms was −0.028 on D1, p=0.29, and 0.059 on D4, p=0.024, bootstrap; effect increased with experience, D4-D1=0.088, p=0.0084, permutation test).

**Figure 2.**
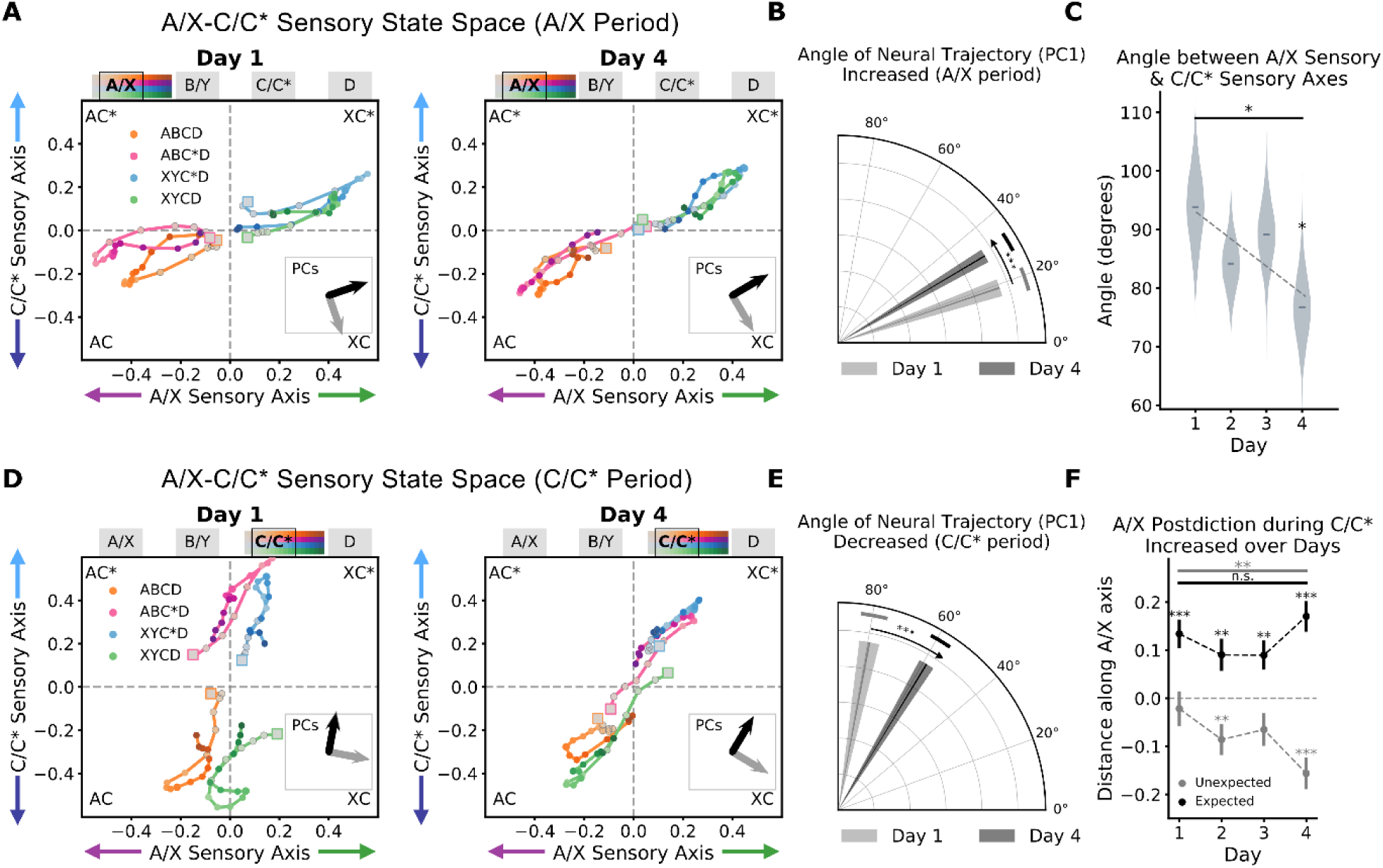
Alignment between A/X and C/C* Representations Facilitates Prediction and Postdiction. **(A)** Neural activity projected into A/X-C/C* state space for Day 1 (left) and Day 4 (right). The x-axis is the magnitude of neural activity projected onto A/X sensory axis; the y-axis is the same neural activity projected onto C/C* sensory axis. Activity is shown from A/X period (−10 to 170 ms). Marker saturation increases with time (key along top). Inset shows PCs of neural trajectories in grey, black arrow size matches percentage of explained variance per PC (plotted in Fig. 3F). **(B)** Increase of first PC angle during A/X period across days 1 and 4. Circular mean and standard deviation of bootstrapped angle shown per day. **(C)** Angle between A/X and C/C* sensory axes decreased across days. Bootstrapped angle shown by violin (stars indicate significant difference from 90 degrees). Horizontal bar marks D4-D1 difference. **(D)** Neural activity during the C/C* stimulus period (340 to 520 ms) projected into A/X-C/C* state space (as in **A**). **(E)** Decrease of first PC angle during C/C* period across days 1 and 4. **(F)** A/X encoding during C/C* period (360-460 ms). Positive values indicate correct A/X encoding. Expected and unexpected trials are shown in black and grey respectively. Note negative A/X values indicate postdiction. For all panels, p-values: * ≤ 0.05, ** ≤ 0.01, *** ≤ 0.001

These results suggest a simple mechanism for making predictions: the neural representations for A/X and C/C* become aligned through experience. This alignment would mean that a sensory input, which moves neural activity along the A/X sensory axis, would also induce movement along the C/C* sensory axis (as seen in Figs. 2A and 2D). To test this, we measured the angle between the A/X sensory axis and the C/C* sensory axis. On Day 1, the angle was 94 degrees, which suggests different stimuli are initially represented by independent, orthogonal representations in auditory cortex. Yet, after experience, the angle significantly decreased, aligning the two representational axes (Fig. 2C; 77 degrees on D4, D4-D1 = −17, p=0.032, permutation test; or with a slope of −4.7 degrees per day, p=0.034, permutation regression test, across days; see SI). These results are consistent with experience forming a common representation for the associated stimuli.

While the alignment of the A/X and C/C* representations facilitates prediction, there is no inherent temporal causality in this mechanism. Therefore, one might expect the C/C* stimulus to also induce a representation of the associated A/X stimulus. Indeed, the C/C* stimulus drove neural activity along the A/X sensory axis (Fig. 1E, black box, and Fig. 1G; D1 = 0.077, p = 0.0008, and D4 = 0.16, p<1/5000, bootstrap test). Again, this effect grew across days (Fig. 1G, D4-D1 = 0.086, p=0.003, permutation test). As with the response to A/X stimulus, the C/C* stimulus also drove neural activity along an angle in the two-dimensional A/X-C/C* state space, encoding the A-C/X-C* association (Fig. 2D). This effect also grew with experience: the angle of trajectories rotated over days, away from primarily encoding C/C* to reflecting the A-C/X-C* association (Fig. 2E, D4-D1 = −20.07 degrees, p=0.0002, permutation test).

This is a postdiction: the representation of a past event is influenced by new incoming information. Postdiction is a common psychological phenomenon thought to facilitate perception; as more information is gathered, our previous beliefs about the world are updated (Eagleman 2000; Vaughn and Eagleman 2013; Aru, Tulver, and Bachmann 2018; Choi and Scholl 2006). Our results suggest prediction and postdiction reflect the same underlying mechanism: they are both explained by alignment of the neural representations underlying associated stimuli. Indeed, the dimensionality of neural activity projected into the A/X-C/C* sensory state space (Fig. 2D) decreased with experience, suggesting there was only a single latent variable (Fig. 3F, during C/C* period, the first PC explained 95% and 98% of the variance in the evoked responses on D1 and D4, both p≤1/5000 against chance by permutation test; D4-D1=3%, p=0.011, permutation test). The formation of a common representation could be due to changes in single neuron selectivity (Sakai and Miyashita 1991) or due to interactions between populations (Fig. S3, see SI).

**Figure 3.**
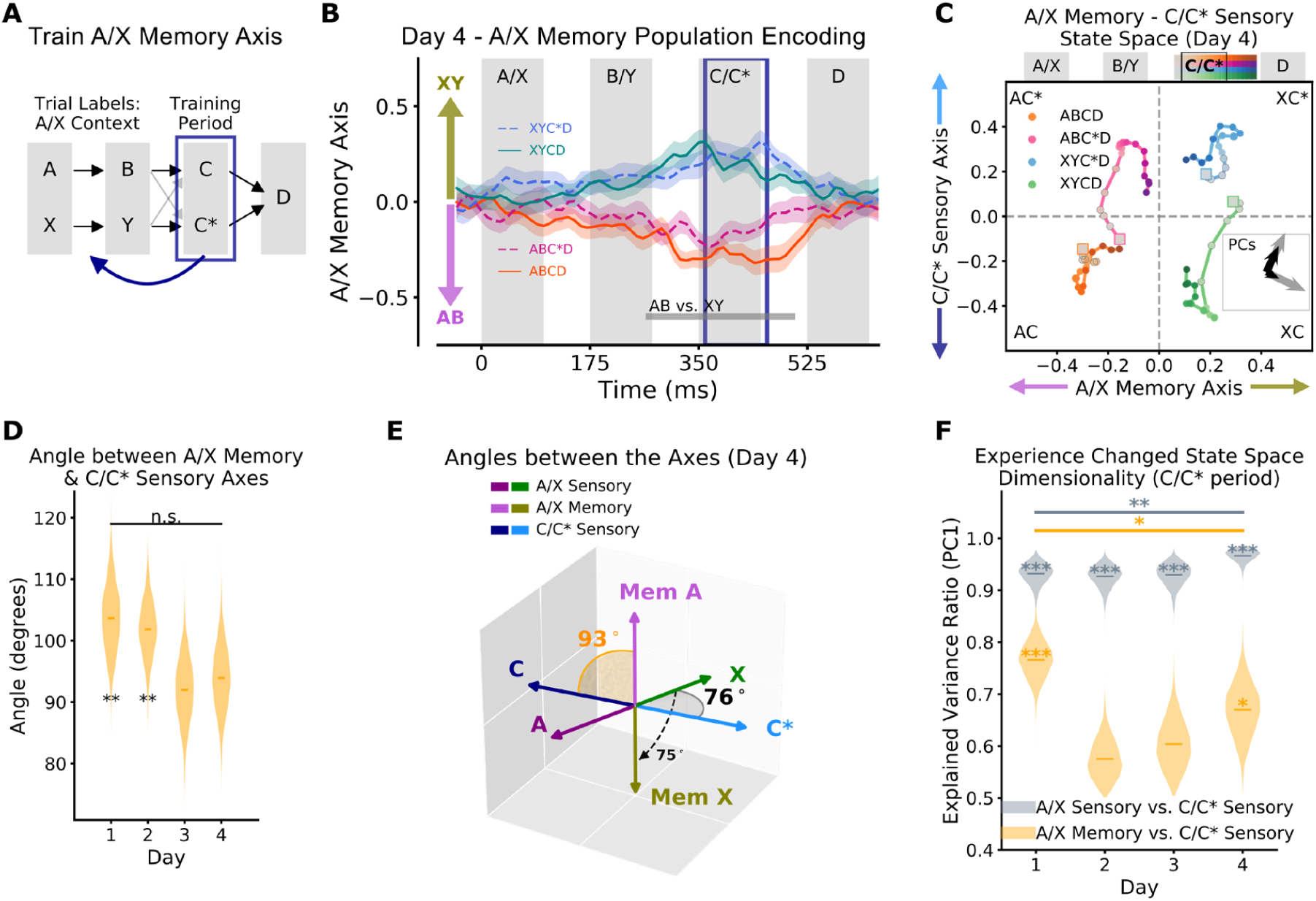
Memory Representations are Orthogonal to Sensory Representations. **(A)** Schematic of training of A/X memory axis. Training used average neural activity taken during C/C* (360-460 ms). Trial types were divided by A/X: (ABCD, ABC*D) vs. (XYCD, XYC*D). **(B)** Neural population encoding of A/X memory information on Day 4. Training period marked by blue box. Positive projections and negative projections indicate XY (green) and AB (purple) memory encoding. Significant differences between AB and XY trials shown by grey bar (p≤0.001, t-test, Bonferroni corrected for multiple comparisons). **(C)** Neural activity projected into A/X memory - C/C* sensory state space on day 4. As in Fig. 2D, but now x-axis is the magnitude of neural activity projected onto A/X memory axis. Activity is shown around C/C* period (340 to 520 ms). **(D)** Angle between A/X memory and C/C* sensory axes. Differences from 90 degrees indicated by stars. **(E)** Schematic showing the angles between the three axes of interest: A/X sensory, C/C* sensory and A/X memory (shown from Day 4). Dashed arrow indicates rotation from the A/X sensory axis to the A/X memory axis. **(F)** Dimensionality of state space around C/C* presentation (340-520 ms). Dimensionality is measured by the percent explained variance (PEV) of the first PC of the neural activity in the A/X sensory – C/C* state space (grey, Fig. 2D) and A/X memory - C/C* state space (orange, Fig. 3C). Violin plots show distribution of bootstrapped PEV. Differences between dimensionality within each state space (grey and yellow) are significant on all days (p≤1/5000, permutation test). For all panels, p-values: * ≤ 0.05, ** ≤ 0.01, *** ≤ 0.001

Yet, because of postdiction, A/X information is lost. Specifically, during unexpected sequences (ABC*D and XYCD), postdiction causes the A/X encoding to reverse, crossing to encode the incorrect context (Fig. 2D, pink and green lines). The impact of postdiction on unexpected stimuli increased with experience (Fig. 2F, A/X unexpected encoding on D1 = −0.022, p = 0.57, bootstrap test; D4 = −0.16, p < 1/5000, bootstrap test; D4-D1 = −0.13, p = 0.003, permutation test; across all days, slope = −0.08, p = 0.007, permutation test). Thus, while sensory axis alignment facilitates prediction (and postdiction), it can also distort the history of sensory representations.

However, it is still important to keep an accurate account of stimulus history. Indeed, short-term memory of recent events is necessary for learning predictions, as it allows for associations across time (Kiyonaga et al. 2017; Summerfield and de Lange 2014). So, we tested whether auditory cortex still maintained A/X information during the C/C* stimulus. Using neural activity during C/C*, we trained a classifier to distinguish the A/X context (Figs. 3A and S2). This defined an ‘A/X memory’ axis, which represented the memory of the A/X context but, surprisingly, did not represent the A/X stimulus itself (Fig. 3B and S4). This suggests the sensory and memory representations were independent. Importantly, A/X information was no longer overwritten by the incoming C/C* stimulus in the two-dimensional A/X memory - C/C* state space (Fig. 3B, C). This was because the A/X memory axis was orthogonal to the C/C* sensory axis (Fig. 3D and 3E; angle=94 degrees on day 4, with a slight, non-significant, decrease towards orthogonality over days, slope = −3.95, p=0.052, permutation regression, D4- D1 = −9.8, p=0.12, permutation test). Furthermore, unlike the A/X – C/C* sensory state space, the dimensionality of the A/X memory – C/C* state space increased with experience (Fig. 3F, explained variance of PC1 during C/C* period on D4-D1 = −10%, p=0.013, permutation test). Together, these results provide evidence for an independent memory representation of context, which avoids interference from the C/C* stimulus.

The importance of both the sensory and memory representations of A/X is reflected by their impact on the representation of C/C* stimuli. On a trial-by-trial basis, the strength of A/X memory encoding (taken 50 ms prior to C/C* onset) was positively correlated with the response to an unexpected C/C* (Fig. S5; on day 4, slope = 0.11, p = 0.01; and non-significantly negatively correlated with expected C/C* encoding strength, slope = −0.03, p = 0.21, bootstrapped linear regression). This is consistent with previous work on predictions that has shown responses to unexpected stimuli are enhanced (while expected are reduced, Fig. 1K, (Dehaene et al. 2015; Kaposvari, Kumar, and Vogels 2018)). Interestingly, we found the opposite relationship between A/X sensory and C/C* encoding, suggesting the two different A/X representations may play different roles in sensory predictions (Fig. S5; expected trials: slope = 0.09, p=0.02, unexpected trials: slope=-0.19, p≤1/5000, bootstrapped linear regression).

Next, we were interested in understanding how A/X context information is transformed from the sensory representation to the memory representation during the sequence. This transformation allows the A/X sensory representation to align with the C/C* representation in order to facilitate prediction, while still maintaining an independent A/X memory axis that is orthogonal to C/C*, and therefore resistant to interference. Two general mechanisms could support this transformation. First, the sensory and memory representations may be represented by independent populations of neurons. This would allow the neurons representing a stimulus to form associations without disrupting the memory representation. Alternatively, the A/X sensory and memory representations may use the same neural population, but change over the course of the sequence, such that the A/X memory and C/C* representations are orthogonal. To distinguish between these hypotheses, we examined how individual neurons encoded the A/X context over time.

Our results show individual neurons in auditory cortex have diverse dynamics. Some neurons maintain their preference for the A/X context throughout the entire trial (Fig. 4A, top) while others change their preference during the trial (Fig. 4A, bottom). To quantify these temporal dynamics across the entire population, we used an unsupervised clustering algorithm (Phenograph, (Levine et al. 2015; Nicosia et al. 2009), see SI). Clustering revealed two broad functional clusters of neurons (see Fig. S6 for d-prime post-hoc clustering validation and cell types along recording arrays). First, ‘stable’ cells maintained their contextual preference across the sequence (Fig. 4B, green and yellow). Second, ‘switching’ neurons switched their A/X contextual preference during the sequence (Fig. 4B, red and blue). Surprisingly, these two broad groups captured the majority of single cell dynamics in auditory cortex (Fig. 4C). Note that the temporal dynamics were not due to neurons having a random preference for the first and second chord in the sequence. The number of stable and switching neurons was greater than would be expected by cells having independent selectivity to each stimulus (Fig. S8, see SI).

**Figure 4.**
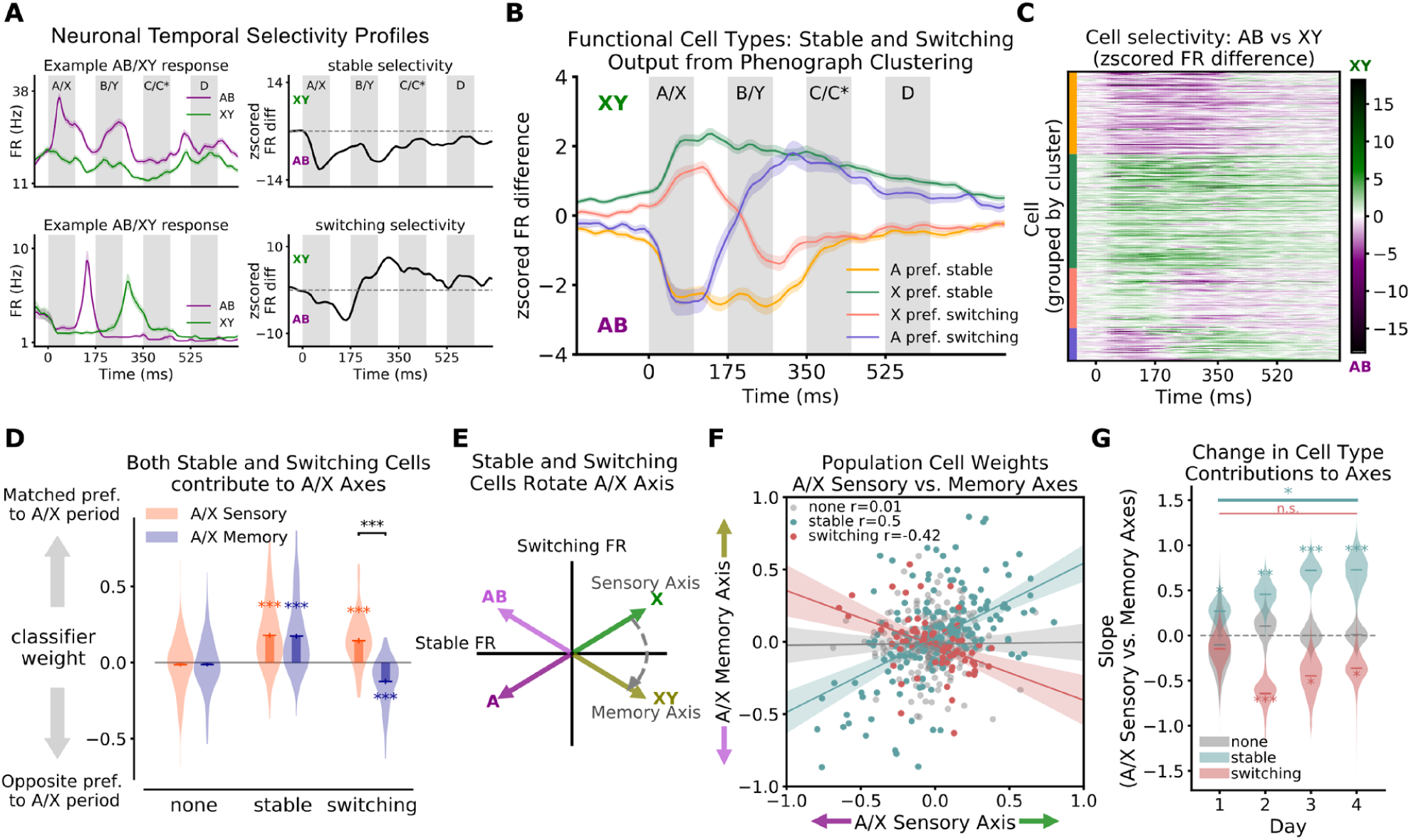
Stable and Switching Neurons Rotate Context Representation from Sensory to Memory. **(A)** A/X selectivity over time for two example neurons. Left column shows the average firing rate in response to A (purple: ABCD, ABC*D) and X (green: XYCD, XYC*D). Right column shows the selectivity of neurons over time, defined as the z-scored firing rate difference of responses to A and X trials (trial types were balanced). **(B)** Unsupervised clustering of selectivity found four clusters. Average selectivity response is shown for each cluster (mean and s.e.m. shown). Purple and pink are switching (32% of neurons; 35% and 65% of which initially preferring AB and XY, respectively, before switching to preferring XY and AB); green and yellow are stable (68% of cells; 42% and 58% preferring AB and XY, respectively). **(C)** Selectivity profiles of the total population (N=522), grouped by cluster. Color saturation indicates z-scored firing rate difference (XY-AB). Colors along y-axis match cluster groups in **B. (D)** Stable and switching cells contribute to both A/X sensory and memory axes. Classifier weights were re-oriented such that positive values indicate a match with preference during A/X period. Cells without significant selectivity at any time (p≤0.025 Bonferroni corrected) were post-hoc removed from clusters (labeled as none), leaving 40% of all neurons in the stable cluster and 13% switching. **(E)** Schematic of how combination of stable and switching cells lead to a rotation of the A/X sensory axis to the A/X memory axis. Stable cells have the same firing rate response to A/X stimuli, during both the A/X and C/C* period (see match between sensory and memory axes). Switching cells reverse their firing rate preference by the memory period. The combination leads to a rotation of encoding space. **(F)** Classifier weights for A/X sensory axis (x-axis) and A/X memory axis (y-axis) for all cells. The lines show the mean and std. of bootstrapped linear regressions for each cell type. **(G)** Experience increases the correlation between the weights of neurons contributing to A/X sensory and memory axes. Slopes (illustrated in **F**) change across days. For all panels, p-values: * ≤ 0.05, ** ≤ 0.01, *** ≤ 0.001

Next, we examined how the stable and switching neuron types contributed to the sensory and memory representations of context. To this end, we examined each cell type’s contribution to each axis (i.e. their classifier weights). Both the stable and switching neurons significantly contributed to the A/X sensory axis (Fig. 4D, orange). Specifically, the A/X sensory axis positively weighted the activity of stable and switching neurons towards their initially preferred stimulus (preference was defined during the presentation of the A or X chord; average weight combined across days for stable neurons = 0.18, for switching = 0.14, both p ≤ 1/5000, bootstrap test). In other words, during the A/X period, both stable and switching neurons increase their activity to their (initially) preferred chord, thereby creating the A/X sensory axis (Fig. 4E).

Similarly, the A/X memory axis relied on the activity of both stable and switching neurons (Fig. 4D, blue). However, while the stable neurons had positive weights (average classifier weight = 0.17, p ≤ 1/5000, bootstrap test), the switching neurons now had negative weights (−0.12, p≤1/5000, bootstrap test; significantly lower than A/X sensory weights, difference = 0.27, p ≤ 1/5000, permutation test). The negative weights of switching neurons reflect their changing preference over the course of the sequence. This makes sense, the neurons’ preference switched over time; thus, the memory axis reflects this new preference (Fig. 4E).

The relationship between the weights of the sensory and memory axes was also evident for individual neurons. The weights of stable neurons were positively correlated across the sensory and memory classifiers (Fig. 4F, green, neurons combined across days slope = 0.5, p<1/5000, bootstrap regression), while the weights of switching neurons were negatively correlated (Fig. 4F, red, slope = −0.42, p<1/5000, bootstrap regression). The difference between stable and switching was significant on all days (D1 = 0.41, p = 0.047; D2 = 1.08, p = 0.0006; D3 = 1.18, p=0.0006; D4 = 1.1, p=0.0004, permutation test). Experience increased the correlations of weights across days (Fig. 4G; D4-D1 slope of stable weights = 0.47, p=0.018; switching = −0.22, p=0.19, permutation test) and increased the difference in weights between the populations (D4-D1 = 0.68, p=0.015, permutation test). Together, these results show both the sensory and memory representations depended on the entire population, arguing against the hypothesis that independent populations represented sensory and memory information.

Our results demonstrate how a combination of stable and switching cell types cause the A/X representation to rotate, away from the sensory representation, to a nearly orthogonal memory representation (Fig. 4E). Thus, the memory representation is orthogonal to the C/C* sensory axis. Now, new stimulus inputs (C/C*), which drive neural activity along the associated A/X sensory axis, do not induce movement along this orthogonal A/X memory axis (Fig. 3C). The dynamics of neural responses we observed in short-term memory representations in mouse auditory cortex are similar to the dynamics found in working memory representations observed in primates (Fig. S4). Indeed, both stable and dynamic representations are seen in monkeys (Murray et al. 2017; Stokes et al. 2013) and their relative contributions to working memory have been debated (Postle 2015; Chaudhuri and Fiete 2016). Our results show how the combination of both response types can transform the sensory representation to a memory representation in a way that reduces interference with future stimuli.

In sum, we have shown a simple neural mechanism underlies prediction: associated representations become aligned over time, such that a contextual stimulus evokes the predicted stimulus representation. This mechanism also leads to postdiction, where previous events are updated based on new stimulus inputs. Finally, we show the neuronal population preserves an accurate history of events by dynamically rotating the information into an orthogonal axis. This dynamic rotation is the result of a combination of stable and dynamic representations at the single neuron level.

## Supplementary Information

### Implicit Learning Paradigm

Mice learned to make predictions in an implicit sequence learning paradigm. On each day, mice were head-fixed and listened to 1500 sequences of four chords (e.g. ABCD); they began naïve to all chords and sequences. Recordings lasted about an hour and were done at the same time each morning (± 1.5 hours).

Within a sequence, each chord lasted 100 ms, and was separated by a 75 ms inter-sound interval (ISI). Inter-trial intervals (ITI) lasted between 500 and 1000 ms (random uniform distribution). Each chord was a combination of 2 frequencies (7/12 of octave apart). Sound waveforms were created in Matlab, with a sample rate of 140 kHz. The frequencies making the chords were drawn from between 10 kHz and 65 kHz. A, B, X, and Y sounds were lower in frequency than C and C* chords. The frequency of D fell between context and C/C* chords. If the frequency of A was less than B, then the frequency of X was greater than Y, and vice versa (8/12 of octave). We used MF1-S speakers (range - 1kHz to 65kHz, Tucker Davis Technologies, Alachua, FL /USA) calibrated with a CM16 microphone (Avisoft-Bioacoustics, Glienicke, Germany) and an Ultramic USB microphone (Dodotronic, Castel Gandolfo RM, Italy). Sound intensity was set to a sound pressure level (SPL) of 70 dB. Sounds were played to left ear. The frequencies and chords were varied across mice.

Each sequence began with a sequence of two sounds, defining one of two contexts. In one context, the A chord was always followed by the B chord (the ‘AB’ context). In the second context, the X chord was always followed by the Y chord (the ‘XY’ context). Context AB was most frequently followed by C (rarely by C*), while context XY was most frequently followed by C* (rarely by C). Expected trials accounted for 68% of trials (e.g. ABCD, XYC*D). Trial counts of ABCD always equaled trial counts of XYC*D. Unexpected trials accounted for 20% of trials (e.g. ABC*D, XYCD). Trial counts of ABC*D always equaled trial counts of XYCD. Importantly, both contexts AB and XY occurred equally per day, as did both predicted stimuli (C and C*). This prevented any *a priori* expectation of any stimulus. The remaining 12% of trials contained an ‘ambiguous’ third stimulus, which was created by combining the frequencies making up the C and C* chords. These trials are not analyzed here. Trial types occurred randomly during the 1500 trials on a given day, according to their probabilities and ensuring equal numbers of trial types, as noted above.

Prior to and after the block of 1500 sequence trials, the C and C* chords were played in isolation for 300 trials in order to measure the stability of representations. As during the sequence, the chords were played for 100 ms. There was a random 500-1000 ms delay between chords.

### Neuronal Recordings

#### Animal Subjects

All animal procedures were approved by the Princeton IACUC and carried out in accordance with National Institute of Health standards. Seven adult male PV cre+/- C57BL6 mice were used for recording and passive learning experiments. Mice were between 13 and 19 weeks old at date of first recording. Animals had free access to food and water and were housed in a reverse light cycle. Experiments were conducted in a sound proofed behavioral chamber.

#### Neural Recordings

Neural activity was recorded at 30kHz using the Intan RHD2000 system (Intan Technologies, Los Angeles, CA). Signals were amplified and digitized on a 32 channel amplifier board (headstage) before being sent (via SPI interface cable) to the USB interface board. Analog signals for speaker were also routed to the interface board, for later alignment of sound timing with neural activity. The Intan recording system software was used to save all data (i.e. sound timing and neural activity).

#### Implant Surgery

While under anesthesia, 32 channel silicon recording arrays (NeuroNexus, Fairfield, CT) were implanted into auditory cortex. Six mice were implanted with a four shank probe (8 electrodes per shank), inserted along the A/P axis. One mouse was implanted with a one shank probe (32 electrodes total). All electrode probes were implanted in right auditory cortex (stereotaxic coordinates from bregma: −2.7 AP and 4.8 ML). Probes were lowered between 970-1400 um in order to target primary auditory cortex (see Fig. S6B for approximate cell locations), although dorsal contacts may have also recorded from secondary auditory cortex. To insert the probe, a dental drill was used to make a small cranial window (typically 3 mm × 3 mm) above the desired coordinates. The silicon probe was lowered slowly, avoiding blood vessels. KwikSil (World Precision Instruments, Sarasota, FL) was used to protect the brain and stabilize the exposed probe. Three screws (miniature self-tapping screws made from #303 stainless steel; J.I. Morris, Oxford, MA) were used to keep the headpost (3D printed at Midwest Prototyping, Blue Mounds, WI) and electrode stable. Ground wires were wrapped around the screw on the opposite side of the brain. Metabond (Parkell, Edgewood, NY) was used to fix all implants to skull.

After surgery, mice were given several days to recover and pain killers (buprenorphine) were given during recovery. Prior to recording sessions, mice were acclimated to handling by the experimenter and head fixation in increments of 15 minutes. Location of silicon probe was confirmed using histology (see Fig. S1C for example electrode placement from one animal). Lesions from electrodes were determined by labeling for astrocytes (GFAP - green).

#### Analysis of Neural Activity

Single units were isolated from the raw 30 kHz signal using Plexon Offline Sorter. Raw data was imported into Plexon Offline Sorter, and filtered using a 350 Hz highpass, 4-pole filter. Next, we applied a common average reference to all channels. Using these traces, we identified clusters of spikes. Animals were excluded from future analysis if they have fewer than 5 single units. We recorded from 10 animals, but only 7 had sufficient single unit activity to be included. From the remaining animals, we found 522 single units across the 4 days of recording.

#### Firing Rate Calculation

The instantaneous firing rate of neurons was estimated at each time point by inverting the inter-spike interval. This trace was smoothed, with a one ms boxcar, and then down sampled to 1000 Hz. Data was then segmented by trial start and end times. For sequence data, trials were taken from 70 ms prior to the A/X chord to 355 ms after the end of the D chord. For the C/C* chord alone, trials were taken to start 70 ms prior to chord onset and end 280 ms after the chord ended. Data was smoothed again with a 20 ms boxcar. All time labels in figures indicate the leading edge of any time frame or window (i.e. including data up to that labeled point). Preprocessing of firing rate data (segmentation and smoothing) was performed in Matlab 2016 (Mathworks, Natick, MA). Also, we computed the z-scored firing rate difference and phenograph clustering in Matlab 2016. All other analyses were performed in Python 3.5.5. For the python analyses (jupyter notebook (Perez and Granger 2007)), we utilized the scipy (Millman and Aivazis 2011; Oliphant 2007), sklearn (Pedregosa et al., n.d.), numpy (Walt, Colbert, and Varoquaux 2011) and pandas (McKinney 2010) packages (specific functions referenced below). All plotting was completed using matplotlib (Hunter 2007).

### Encoding Axis (Classifier) Training

All classifiers were trained using the same procedure. The only difference between classifiers were training period and condition groupings. For a given comparison, we trained one classifier for each mouse, on a given day.

#### Feature Selection

The feature vector for each trial was the averaged firing rate response during sound presentation periods, shifted forward by 10 ms to account for onset time of sensory response (i.e. 10 to 110ms after stimulus onset). For the cross temporal decoding, we trained multiple classifiers across time, in a sliding window fashion. For these sets of classifiers, we used average firing rates in 25 ms time bins, moved by 10 ms across the sequence.

#### Classifier Training Labels

The classifiers were trained to distinguish between groups of conditions (e.g. AB trials, labeled as 0, versus XY trials, labeled as 1). All classifiers trained, and their respective comparisons, are listed below. Generally, classifiers fell into one of two categories: sensory classifiers distinguished the two possible sounds at the time that they occurred within the sequence and the memory classifier distinguished the two possible contexts during the latter part of the trial, after the chords had occurred.

#### List of Classifiers Trained

*\**(conditions balanced in both training and testing)

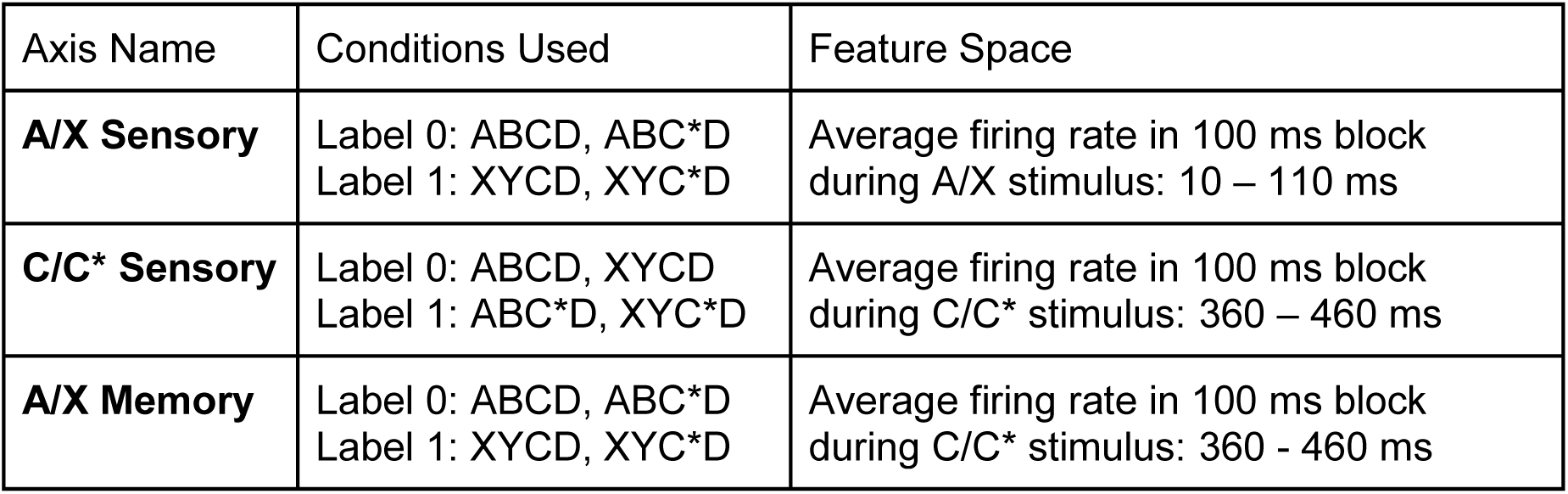

#### Classifier Type

The classifier was a linear classifier trained via stochastic gradient descent with a hinge loss function, which is equivalent to a standard SVM (support vector machine). To train the classifier we used the stochastic gradient descent SGDClassifier function in the sklearn.linear_model package (sklearn version 0.19)(Pedregosa et al., n.d.) for Python3 (version 3.5.5).

#### Classifier Regularization

To minimize overfitting of the classifiers, we used an elastic net regularization to increase sparsity of the weight vector and to minimize the number of non-zero weights. Parameters were the same for all classifiers: L1 ratio (the elastic net parameter specifying the ratio of L1 to L2 penalties) was set to 0.65, alpha (the regularization amount) was set to 0.01, the learning rate was set to 0.00001, and the number of iterations was set to 1000. See the SGDClassifier function for details.

#### Cross-Validated Testing of the Classifier

Prior to training, 10% of trials were withheld as a test set to allow for cross-validated testing of the classifier. All figures using classifiers were created only with this withheld test data. Both the training and test dataset contained equal counts of all conditions, regardless of how conditions were combined for a given classification. For example, for classifying AB vs. XY trials, there were equal counts of ABCD, ABC*D, XYCD and XYC*D in the dataset (note: this is different from what the animal experienced but avoids bias in the classifier). For each comparison, 100 shuffles of the training data were performed to ensure all trials were included in training the classifier. The classifier was calculated by taking the mean (intercept and weights) of these 100 trained classifiers.

#### Classification Accuracy of the Classifiers

Once trained, the classifier defines a hyperplane in feature space (firing rate of neurons). The distance of population activity from this hyperplane may be used to label each sample as a member of one condition or another (e.g. context AB or XY). The label maybe be correct (True positive or True negative) or incorrect (False positive or False negative). To determine classifier accuracy, we calculated the area under the curve (AUC) of the receiver operator characteristic (ROC) of the true positive rate verses the false positive rate as the decision boundary is varied (Fig. S2). To do this, we used the roc_curve and auc functions in the sklearn.metrics package.

### Projection onto Encoding Axis

#### Projection onto 1D Encoding Axis Defined by a Classifier

Once trained, a linear classifier is defined by a vector, normal to the separating hyperplane and an intercept. The distance of a given sample from that hyperplane is the dot product of that vector with the sample (e.g. a trial firing rate across N neurons), plus the intercept. Thus, the classifier may be used as a 1D encoding axis, by which the N-Dimensional neural activity may be projected to determine its encoding of a given condition on a given trial (Fig. 1D). The sample’s distance from hyperplane reflects its similarity with each label used to train the classifier.

To calculate population encoding, we took data from each mouse, and projected the full neural population activity onto the 1D axis defined by the trained classifier for that mouse. Specifically, for each trial, at each time point (25 ms bin), we took the vector of firing rates across all neurons and projected it onto the encoding axis of interest. This value was then z-scored across conditions (i.e. subtracted mean and divided by standard deviation, calculated from all conditions). This z-score captures the relative separation between conditions across time and ignores any absolute drift in firing rates occurring over time (essentially allowing the intercept of the classifier to change over time, which is important to capture relative changes in the firing rate across the population). Furthermore, the z-score allows us to combine responses across mice. The projection is signed such that a positive value indicates strong encoding of the positively labeled condition; likewise, negative values indicate strong encoding of the negatively labeled condition. For example, for the classifier trained to distinguish AB trials (label 0) from XY trials (label 1), if the population activity had a positive distance from the hyperplane, then it is representing the context XY. In contrast, a negative distance from the A/X hyperplane would represent context AB. Figure 1E shows the A/X population encoding over the sequence on days 1 and 4. Figure 1H shows the C/C* population encoding over the sequence on days 1 and 4. To examine how encoding of context (AB vs. XY) and C/C* stimulus changes over time, we performed t-tests (function: ttest_ind from scipy.stats package) on each time bin. The neural population was said to be carrying significant information about a stimulus (or memory) if the associated p-value was less than or equal to 0.001, bonferroni corrected for multiple comparisons across time (note: all tests were done on withheld test data, as described above).

#### Distance from Axis: Encoding Strength / Accuracy

We also tested encoding accuracy during a given time frame (e.g. Fig. 1F, G, I, J, Fig. 2F, and Fig. S4). To calculate absolute encoding strength (i.e. combined across conditions), we flipped the sign of trials with a negative label and calculated the mean distance across trials. For example, when testing A/X encoding (Fig. 1F, J), we flipped the sign of trials of A conditions: ABCD and ABC*D. Likewise, when testing C/C* encoding (Fig. 1G, I), we flipped the sign of trials of C conditions: ABCD and XYCD. Then we combined across all trials and reported the average distance from the hyperplane.

To test significance of encoding strength during a time period, we estimated the distribution of the mean encoding strength with a bootstrap procedure (5000 samples). The bootstrap sampled randomly from the distribution of observed values (with replacement, (Manly 1997). The resulting distribution allowed us to estimate the probability of observing an average encoding value less than or equal to zero (this is reported as our p-value). Furthermore, variability in the linear regression (function: linregress from package scipy.stats) across days was estimated using these bootstrapped values. To test for pairwise differences in encoding strength across days, we performed a permutation test on the difference of mean encoding on each day (Manly 1997). We shuffled the day labels 4999 times (adding the original observation to make 5000 permutations), generating a null distribution of differences across days. Comparing the observed difference to this null distribution allowed us to estimate the p-value.

#### Prediction prior to onset of C/C* Stimulus

As described in the main text, we found evidence of A-C and X-C* associations along the respective A/X and C/C* sensory axes during each stimulus presentation (A/X period Fig. 1J, and C/C* period Fig. 1G). In addition, we found evidence for predictions prior to the onset of the C/C* stimulus (Fig 1; Fig 2D). In the 50 ms before the onset of the C/C* stimulus, the A/X sensory axis carried the memory of the A/X stimulus (D1=0.092, p=0.0008, bootstrap; D4=0.077, p=0.004, bootstrap test). On day 4, the C/C* sensory axis already encoded the predicted stimulus (Fig. 2D, right). If the animal previously heard A (pink and orange lines) then, prior to the onset of C/C*, neural activity was biased in the C direction (and vice-versa for X and C*; AC/XC* prediction = 0.059 on D4, p=0.024, bootstrap). Again, experience increased this prediction; it was not significant on Day 1 (−0.028, p=0.29, bootstrap) and increased over days (D4-D1=0.088, p=0.0084, permutation test).

#### Calculation of Angle between Axes (Trained Hyperplanes)

To examine how experience influences representations, we measured the angle between encoding axes. Classifier weights were normalized to a length of 1 for each animal and then concatenated across animals to form a single vector. We calculated the angle between a pair of hyperplanes by inverting the dot product of each hyperplane’s normal vector:

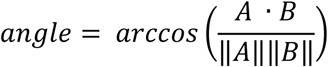

#### Neural Mechanism Aligning A/X and C/C* Sensory Representations

Our results show the A/X sensory and C/C* sensory representations become aligned with experience. This likely happens due to changes in the selectivity of individual neurons, such that neurons become responsive to both associated stimuli (e.g. neurons, selectively responsive to A, become responsive to C as well). There are multiple mechanisms inducing these changes in selectivity (schematized in Fig. S3). First, the connectivity of inputs to auditory cortex could change such that A/X and C/C* neurons respond to the presentation of either stimulus (Fig. S3A). Second, unidirectional lateral connections could facilitate a transformation from the A/X representation to the C/C* representation (Fig. S3B). If A/X neurons increase their connection to C/C* cells, then the onset of the A/X stimulus will lead to an initial response from the A/X neurons, followed by a delayed response of the C/C* neurons. Third, bidirectional lateral connections could facilitate a transformation either from the A/X representation to the C/C* representation, during the presentation of A/X, or from the C/C* representation to the A/X representation, during the presentation of C/C* (Fig. S3C).

Importantly, all three potential mechanisms would lead to a conjoined representation within auditory cortex. This would confuse the information available to downstream brain regions. By training linear classifiers, we captured how a simple, linear downstream neuron would decode information from the representation in auditory cortex. As we show in the main manuscript, the conjoining of representations leads to both prediction (Fig. 1J) and postdiction (Fig. 1G, Fig. 2F).

One way to discriminate these three models is to examine how A/X and C/C* information evolves in response to the A/X and C/C* stimulus. If the selectivity of individual neurons is changing (the first hypothesis, Fig. S3A), then there should be no timing difference between the A/X and C/C* responses to either the A/X or C/C* stimulus. In contrast, if unidirectional lateral connections form between A/X and C/C* neurons (second hypothesis, Fig. S3B), then C/C* representations should follow the A/X representations during the presentation of A/X alone (due to the asymmetric connection from A/X to C/C*). Finally, if bidirectional lateral connections are strengthened between A/X and C/C* neurons (third hypothesis, Fig. S3C), then there should be a timing delay between the A/X representation and the C/C* representation, during both A/X and C/C* stimulus presentations.

To examine the timing of A/X sensory and C/C* sensory encoding, we calculated the time to reach significant sensory encoding during each time period (A/X and C/C*) along each axis. For a given window of interest (e.g. around presentation of the A/X stimulus, Day 1 shown in Fig. S3D, Day 4 shown in Fig. S3F), we estimated the encoding strength over time (interpolating to get higher temporal precision). Significance was taken as the time to reach statistical threshold (p ≤ .001, Bonferroni corrected). On day 1 and 4, we found that during A/X period, information is first represented along the A/X axis (D1=17 ms, D4=24 ms) and then on the C/C* axis (D1, Fig. S3D, 44 ms, lag=26.5 ms, p≤1/5000, D4, Fig. S3F, 42 ms, lag=17.8 ms, p=0.0002). In contrast, during the C/C*period, information is first represented along the C/C* axis and then along the A/X axis (Fig. S3E, D1=354 ms and 406 ms for C/C* and A/X respectively, lag = 51.4 ms, p=0.0002; Fig. S3G; D4=377 ms and 396 ms for C/C* and A/X, respectively; lag=18.9ms, p=0.0056). These timing results suggest A/X and C/C* representations became more similar through learning in bidirectional lateral connections (Fig. S3C).

Furthermore, we tested if there were significant differences in the percentages of cells in the A/X and C/C* sensory axes. We combined classifiers across animals by normalizing the weight vector for each animal and then concatenating the vectors. To measure whether classifiers relied on the same proportion of neurons, we systematically varied the weight threshold for a neuron to be defined as ‘contributing’ to a classifier. For each threshold, we calculated the percent of neurons with classifier weights above this threshold for both the A/X and C/C* sensory axes. Plotting these against each other yields a ROC-like curve (Fig. S3H). If one classifier utilized a higher percentage of cells than the other, the area under the curve (AUC) would be significantly different than .5. Across all four days, using a bootstrap (subsampling neurons 5000 times), we failed to find AUCs significantly different than .5, suggesting an equal portion of cells were involved in each sensory classifier (Fig. S3I, D1 p=0.13, D2 p=0.35, D3 p=0.46, D4 p=0.30).

These results argue against a unidirectional learning effect (which would lead to more neurons involved in the A/X representation than the C/C* representation). Altogether, these results suggest there is a recurrent dynamic between both sensory encodings, which may underlie the alignment between representations.

### Dimensionality and Angle of Neural Activity in 2D State Spaces

#### Calculation of Principal Components and their Explained Variance

To understand the dimensionality of the temporal trajectories, we calculated the principle components (PCs) of neural activity in the 2D state spaces using the PCA function in the sklearn.decomposition package (Fig. 2A (inset), 2B, 2D (inset), E, and 3C (inset)). The two-dimensional state spaces are created by combining the individual encoding axes. Neural activity from every 25 ms was projected onto both encoding axes and averaged across trials. Data was combined from all four conditions (ABCD, ABC*D, XYC*D, XYCD) and the PCs were calculated. In this way, the PCs capture the direction of neural activity dynamics within the state space.

The angle of the first PC in the state space captures the information represented by neural activity. Specifically, if the neural activity moves only horizontal or vertical than the neural activity is only encoding that one type of information. To capture the increase in encoding along both the A/X and C/C* sensory axes, we calculated the PC angle during both the A/X period and C/C* period. During the A/X period, over days of experience, the PC angle increases from horizontal (i.e. only A/X encoding) to diagonal (i.e. both A/X and C/C* encoding; Fig. 2B). This indicates an increase in the A-C/X-C* association during A/X. Similarly, during the C/C* period, over days of experience, the PC angle decreased from vertical, also demonstrating the association (Fig. 2E).

The ratio of the percent of variance captured by each PC is proportional to the dimensionality of the representation. For example, if activity flows along a single dimension, such as in the A/X-C/C* sensory state space (Fig. 2D), then the first PC will explain nearly all of the variance in the neural activity over time. In contrast, if this ratio decreases such that the first two PCs explain similar amounts of variance (e.g. in the A/X memory – C/C* sensory state space, Fig. 3C), then this would indicate neural activity explores the full two-dimensional space.

##### Bootstrapped Distributions

We used a bootstrap process to estimate the distribution of PC angle and the ratio of explained variance of the PCs. The trials used to estimate each timepoint of neural activity were sampled randomly (with replacement) and PCs were recalculated. This process was repeated 5000 times to estimate the distribution of each variable. These distributions were then used for plotting or for calculating linear regressions on the data across days. When used for linear regression, a line was fit to a single sample drawn from the bootstrap on each day (resulting in 5000 fits). This process allowed us to estimate the distribution of slopes and intercepts.

##### Permutation Tests

In order to determine if the observed dimensionality was lower than expected by chance (Fig. 3F), we created a null distribution of projections into the 2D state space by randomly permuting the time labels of each point, separately in both the x and y dimensions. In this way, we kept the distribution of raw activity values, but broke their temporal association. We recalculated the PCs and the ratio of explained variance for each permuted data set. This process was repeated 4999 times (the 5000th data point is the unpermutated, observed data) in order to estimate the probability of observing the original percent explained variance ratio by chance.

To calculate differences across state spaces (across days or state spaces), we shuffled points between the two state spaces in order to maintain the distribution of activity, but break the association to a specific state space. For example, to compare the neural trajectories between days 1 and 4, we shuffled the day labels on each data point, and then calculated the difference in PCs, angles, and percent explained variance. We performed 4999 shuffles (5000^th^ point is unpermutated data) and used this distribution to calculate the probability of observing our original data.

### A/X Memory Axis Compared to A/X Sensory Axis

As described in the main manuscript, A/X information is lost along the A/X sensory axis when an unexpected C/C* stimulus is presented (Fig. 2). To test if A/X information persisted during the C/C* stimulus, we trained a classifier during the C/C* period (360 to 460 ms) on distinguishing A trials (ABCD, ABC*D) from X trials (XYCD, XYC*D; Fig. 3A). Thus, the main difference between the A/X sensory and memory classifier was the time period of neural activity used for training (A/X sensory=10 to 110 ms, i.e. during the presentation of A/X, A/X memory=360 to 460ms, i.e. during the presentation of C/C*). The A/X memory classifier performed above chance (Fig. S2D). To examine how A/X memory information evolved over the course of the sequence, we projected neural population firing activity (as described above) on day 1 and day 4. Again, there were significant differences between A/X context trials around the presentation of C/C* (Fig. S4A and B, grey bars p ≤ .001, Bonferroni corrected t-test). In the A/X memory - C/C* sensory state space the memory of A/X stimuli is preserved during the presentation of C/C* on both day 1 and day 4 (Fig. S4, C and D, respectively). This is reflected by the fact that the dimensionality of activity in the A/X memory - C/C* sensory state space was significantly greater than the A/X sensory - C/C* sensory state space (Fig. 3F; note: the neural activity is the same in both cases, just projected differently, D1 diff = 18%, D2 diff = 40%, D3 diff = 38%, D4 diff = 31%, p≤1/5000 for all days).

To understand how A/X encoding evolved over time, we compared A/X encoding along both sensory and memory axes at each stimulus period in the sequence (A/X, B/Y, and C/C* in Fig. S4E, F and G respectively). During A/X presentation, A/X encoding was stronger on the A/X sensory axis. During the B/Y presentation, A/X encoding was weak along both sensory and memory axes. Lastly, during the C/C* presentation, A/X encoding was stronger on the A/X memory axis, compared to the A/X sensory axis. In order to gain a full picture of the evolving A/X axis, we performed cross temporal classification. We trained classifiers on 25 ms time bins throughout the sequence to distinguish between A and X conditions (Fig. S4H). The A/X classifiers performed well around the time of training, but only generalized locally in time. These results suggest a dynamic population coding of A/X information that evolves over time. As detailed in the main manuscript, we find that individual neurons show similar dynamics – some have a stable preference for A/X stimuli across time, while others switch their preference during the sequence. The combination of both stable and switching representations is what determines how the population representation evolves during the sequence.

### The Strength of A/X Sensory and Memory Encoding Impacts C/C* Encoding

To test the impact of context (A/X) encoding on perception, we tested, on a trial by trial basis, whether the strength of context was correlated with the strength of C/C* stimulus encoding. Based on previous work, we anticipated contextual information would impact sensory processing in one of two different ways. First, in the predictive coding framework (Rao and Ballard 1999), contextual information could set an expectation for the upcoming stimulus. In this framework, unexpected stimuli evoke a stronger response (due to a prediction error). This is consistent with our observations (Fig. 1H, I). Therefore, one might expect that stronger contextual information on a given trial could lead to a stronger response to unexpected stimuli on that trial. In other words, there should be a positive correlation between context and unexpected stimuli. Second, the representation of contextual information could predict future stimuli, such that the expected stimulus is enhanced (i.e. it would facilitate the sensory response). In this case, one would predict a positive correlation between the strength of context encoding and the strength of the representation of the expected stimulus.

To test these two hypotheses, we examined the correlation between the strength of A/X context encoding on a given trial with the strength of the representation of either expected or unexpected C/C* stimuli on day 4. For each trial, neural activity was averaged from the 50 ms before the onset of the C/C* stimulus (300-350 ms after the onset of the A stimulus). This activity was then projected onto either the A/X sensory axis or A/X memory axis (as described above). Encoding strength was taken as the magnitude of activity in the correct direction along the axis (i.e. positive values indicate correct encoding and negative values indicate incorrect encoding). To examine expected and unexpected stimuli independently, trials were sorted by whether the stimuli were expected for the given context (i.e. ABCD and XYC*D trials were expected while ABC*D and XYCD trials were unexpected). The strength of context and C/C* representation was then correlated by linear regression (function: linregress from package scipy.stats). As with other analyses, we bootstrapped the linear regression fit by sampling trial data with replacement (5000 repetitions).

We found encoding along the two A/X axes, sensory and memory, had opposing influences on C/C* representation. First, the strength of the A/X sensory representation had a facilitating effect. On expected trials A/X sensory encoding strength was positively correlated with C/C* sensory encoding strength across trials (Fig. S5A, slope=0.086, p=0.019, bootstrapped linear regression). On unexpected trials, A/X sensory encoding strength was negatively correlated with C/C* sensory encoding strength (Fig. S5B, slope=-0.19, p<1/5000, bootstrapped linear regression). These results are consistent with the alignment of A/X sensory and C/C* sensory representation: A/X sensory representations facilitate the predicted response, while also interfering with the unexpected response.

Second, the A/X memory representation had the opposite effect on the C/C* response. On unexpected trials, the strength of A/X memory encoding was positively correlated with C/C* sensory encoding strength across trials (Fig. S5D, slope=0.11, p=0.011, bootstrapped linear regression). On expected trials, the strength of A/X memory encoding was weakly (non-significantly) negatively correlated with C/C* sensory encoding (Fig. S5C slope=-0.033, p=0.2). These results are consistent with a predictive coding framework, which hypothesizes the response to expected stimuli should be reduced (as they are explained away), while unexpected stimuli should be enhanced (as there is a prediction error). Taken together, these results suggest that both A/X sensory and memory axes significantly impact sensory processing, but may play different roles in facilitating sensory responses or making predictions.

### Temporal Selectivity Profiles

As described in the main manuscript, we found the population encoding of A/X sensory and memory representations were largely independent. To understand how these different representations evolved during the trial, we examined how contextual information was represented by individual neurons. To this end, we calculated how each neuron represented context over time. Selectivity was measured as the difference in firing rate for the AB context and the XY context. As before, the trial counts for each sequence were balanced, ensuring the neurons were not responding to the C/C* stimulus. The difference in firing rate between contexts was calculated in 25 ms time bins, shifted by 10 ms over the entire trial (from −160 ms to 790 ms, relative to the onset of the A/X stimulus). To normalize the firing rate difference, we z-scored the observed difference in firing rate by a null distribution, which was created by randomly permuting the trial labels (i.e. shuffling the responses to AB and XY contexts). Using these shuffles, we z-scored the firing rate difference using the following equation.

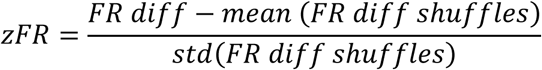

This resulted in a z-scored measure of contextual selectivity for each neuron across the entire time period of the trial (examples in Fig. 4A).

### Clustering of Neuron Selectivity into Functional Cell Types

#### Phenograph

To test whether the timecourse of selectivity of neurons were clustered, we used the unsupervised ‘Phenograph’ clustering algorithm (Nicosia et al. 2009) to cluster the z-scored selectivity profiles (created as above). Briefly, the Phenograph algorithm works by constructing a directed graph of data points, where a given data point is connected to its *k* closest neighbors. Distance between data points was measured as their Euclidean distance in the 96 dimensional space of the full temporal profile of selectivity. Following previous work (Levine et al. 2015), we used *k* = 40. Once this directed graph is constructed, Phenograph uses the Louvain clustering algorithm to cluster data points into groups (Blondel et al. 2008). Note that this algorithm is unsupervised; the only parameter set is the number of local connections (*k*), although the algorithm is robust to changes in this parameter (Levine et al. 2015). The Phenograph algorithm identified 4 clusters of temporal selectivity (Fig. 4B).

#### Phenograph Validation: D-prime

To validate the resulting clusters (outside the Phenograph algorithm), we calculated a sensitivity index between each pair of clusters (Figure S6A). The sensitivity index, often referred to as d-prime, measures the distance between clusters. For each cluster being compared, the distance between cluster means is calculated (using Euclidean distance in the full 96-dimensional space). This is divided by the square root of the average of the variances within each cluster:

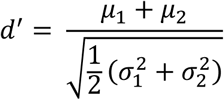

Where 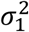 and 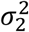 are the variances and *μ*_1_ and *μ*_2_ are the means of the two clusters being compared. To test whether the observed d-primes were significantly greater than expected by chance, we performed permutation tests on each paired d-prime calculation. For each pairwise cluster comparison, we shuffled the labels of each cluster (999 permutations), recalculated d-prime and added the unpermutated d-prime calculation to distribution. Using this null distribution, we calculated the probability of each observed d-prime. All pairwise cluster comparisons were significantly separated (p≤1/1000; Figure S6A).

#### Spatial Clustering of Functional Clusters

To test whether there was anatomical clustering of the observed functional clusters, we estimated each neuron’s anatomical location based on the location of the implanted electrodes. We did not see clear spatial clustering of the functional clusters of neurons (Fig. S6B). The functional clusters were intermingled, suggesting that the different neuron types were not due to differences in recording location.

#### Estimating the Classifier Weights for each Functional Cluster Type

Next, we were interested in understanding the relationship between the functionally-defined cell types from our clustering and the population representations of context. Specifically, we wanted to test if both functional neuron types (stable, switching) were involved in both the A/X sensory and memory axes (Fig. 4). First, to mitigate noise, we post-hoc identified a ‘none’ cluster, in which the selectivity failed to reach statistical significance over the sequence time course (p<=.025, Bonferroni corrected). Then, we averaged the classifier weights for a given encoding axis (e.g. A/X sensory) for each functional cluster (Fig. 4D). Each functional cluster (stable and switching) has two subgroups: neurons that initially preferred the A stimulus (i.e. the yellow and purple lines in Fig. 4B) and those that initially preferred the X stimulus (i.e. the pink and green lines in Fig. 4B). In order to combine weights from both A-preferring and X-preferring neurons, the weights of all neurons that initially preferred the A stimulus were flipped by multiplying by −1. Therefore, the weights presented in Figure 4B are measured with respect to the neuron’s initial preferred direction. As before, classifier weights were length normalized within each animal before combining across animals, to avoid over-weighting animals with more neurons. Then, all weights were averaged together for each of the two resulting functional cell types (stable and switching neurons from all days). The resulting average weights, for each A/X axis (sensory and memory), are shown in Figure 4D. Initially, for the A/X sensory axis, both stable and switching neurons represent their preferred stimulus (either A or X). This is by definition: by flipping the weights of A-preferring neurons, we ensure all weights are positive during the beginning of the sequence (the A/X presentation). Importantly, the classifier weights for the A/X memory axis were positive for the stable neurons, but reversed for the switching neurons (to negative weights). This is consistent with the switching neurons changing their selectivity during the sequence.

It is important to note that the combination of stable and switching neurons is what changes the representation of A/X during the sequence. If all the neuron were stable, the sensory and memory axes would be the same. Likewise, if all the neurons switched their selectivity, the sensory and memory axes would just reverse the direction of the encoding axis. Instead, as seen in Figure 4E, it is the combination of both stable and switching neurons that allows the memory encoding axes to rotate away from the sensory encoding axes.

#### Estimating the Classifier Weights for Individual Neurons

Next, we tested whether the classifier weights of individual neurons changed between the sensory and memory encoding. To test this, we measured the correlation between an individual neuron’s contribution to the A/X sensory classifier and its contribution to the A/X memory classifier (i.e. the correlation between classifier weights). Similar to above, we labeled each neuron by its functional cluster (stable, switching, or none) and length-normalized the weight vector within each animal, before combining across all animals and all days. We plotted each neurons weight in the context sensory and memory axes, organized by their functionally-defined cluster (Fig 4F; x-axis: A/X sensory axis, y-axis: A/X memory axis). We quantified the relationship by calculating linear regressions for each functional cluster. Statistical significance of each linear regression (scipy.stats.linregress) was determined by using a bootstrap. We subsampled the population of neurons (for each function cluster) 5000 times, each time fitting the linear regression. The fitted lines and the standard deviation of the bootstrap can be viewed in Fig. 4F. As with the full population, the weights of individual ‘stable’ neurons were positively correlated across the sensory and memory classifiers, suggesting they maintain their role (and that strongly encoding neurons contributed well to both). In contrast, the weights of ‘switching’ neurons were negatively correlated between the classifiers, suggesting individual neurons dynamically change their contribution to the sensory and memory representations.

#### Classifier Weights Change with Experience

To test how experience impacted the correlation of classifier weights, we recalculated these regressions on each day. The distribution of slopes per day are illustrated by violin plots in Figure 4G. On each day, using the bootstrapped slopes, we also tested if stable and switching cells had positive and negative slopes respectively (Fig. 4G, stars indicated significance). To test if experience changed the correlation between day 1 and day 4, we took the difference of the slopes calculated on each day. To estimate the likelihood of observing this difference, we compared the observed difference in slope to the difference in slope when the day labels were randomly shuffled (4999 shuffles, where the 5000^th^ was calculated using unpermutated data). We found a positive correlation in stable cell contribution to A/X sensory axes on each day (D1=0.3, p=0.015, D2=0.44, p=0.002, D3=0.67, p<1/5000, D4=0.59, p=0.0004). This correlation increased between day 1 and 4 (Fig. 4 horizontal line, D4-D1=0.47, p=0.013). While the negative correlation in switching cell contributions to each axis was not significant on day 1 (D1=-0.15, p=0.18), it was significantly negative on the rest of days (D2=-0.66, p=0.0004, D3=- 0.63, p=0.018, D4=-0.44, p=0.032). However, the change across days was not significant (D4-D1=-0.22, p=0.2).

We also examined if correlations across functional groups were significantly different. To test for significant differences, we shuffled functional labels and then recalculated the correlation for each group and the difference between groups (4999 permutations, 5000^th^ was difference of unpermutated data). Using this distribution, we determined differences between the slopes of switching and stable cells on each day (D1 diff. = 0.41, p=0.046, D2 diff. = 1.08, p=0.0004, D3 diff. = 1.18, p=0.0018, D4 diff. = 1.1 p=0.0004). This difference significantly changed over days (D4-D1 = 0.68, p=0.018). Significance of the change across days was determined with a permutation test; we shuffled the day label within each functional group, then calculated the day difference, and finally computed the difference across days (4999 permutations, 5000^th^ was difference of unpermutated data).

### Excluding Alternative Explanations for Changes in Context Preference

As noted above, we observed two functionally-defined classes of neurons: those with stable contextual (A/X) preference and those that switch their preference. However, one concern is that these functional classes simply reflect the fact that neurons have different patterns of selectivity for the A/X and B/Y stimuli. For example, a ‘switching’ neuron could simply prefer A over X and then Y over B. To test this hypothesis, we built, fit, and compared models to explain the observed cell selectivity in the first two stimulus periods (A/X and B/Y) of the sequence.

#### Independent Model

The ‘Independent’ model, assumed cells had random, independent selectivity at each stimulus period (Fig. S7A; columns indicate A/X period response; rows indicated B/Y period response). More specifically, a cell might have some probability of preferring A or X and then either B or Y. In this scenario, where each selectivity preference is independent from the next, the probability of a cell preferring A then B, is the product of the two probabilities (*pAB* = *pA* * *pB*). Additionally, not all cells are necessarily responsive to stimuli. To capture this, the model allowed some neurons to be unresponsive (with probability *pNR*), while the remaining are responsive (with probability *pR* = 1 – *pNR*). Therefore, altogether, the probability of observing A selectivity, followed by B selectivity can be written as:

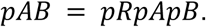

Similar equations can be written for the probability of observing AY (*pAY* = *pRpApY*), XB (*pXB* = *pRpXpB*), and XY (*pXY* = *pRpXpY*). At any given period, a cell may be responsive (*pR*), but not selective for either presented stimulus. For example, a responsive cell may prefer neither A or X with some probability (*p*0_1_=1 - *pA* - *pX*). Given this, we can determine the probability of observing no selectivity during the initial period (A/X), followed by B or Y selectivity:

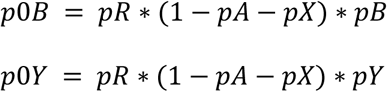

Likewise, we can write the probabilities of observing selectivity, during the initial period (A/X), but no selectivity during B/Y (*p*0_2_=1 - *pB* - *pY*). Here, we can write the probability of observing A0 or X0 as:

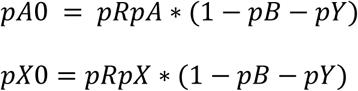

Finally, there are two ways, in which a cell exhibits no response during both the A/X and B/Y stimulus (i.e. 00). First, the cell my not be responsive (*pNR*). Second, the cell may be responsive, but selective for none of the stimuli. Combining, we can write the probability of observing no selectivity as:

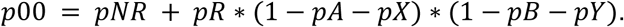

The entire table of probabilities for the independent model is included in Figure S7A. This full model assumes the likelihood of selectivity is independent for each stimulus. However, *a priori*, there is no reason why these likelihoods will be different and so we developed a simpler alternative model (the ‘Simplified Independent’ model) where *pA* = *pB* = *pX* = *pY* = *pS* (Fig. S7B).

#### Stable & Switching Model

Alternatively, the functional classes may reflect the existence of two groups of cells which have stable or switching preferences for the contextual (AB/XY) stimuli. Indeed, this is consistent with the observation that many cells have significant preference beyond the B/Y period, as indicated by their weights in the A/X memory classifier. To test this against the Independent model, we designed a ‘Stable & Switching’ model, in which neurons can have a single preference (AB or XY) and either maintain or change this preference over time. To model this, we defined the probability of a neuron having a stable preference as *pSt* and the probability of switching preferences as *ppp*. Remaining cells were not part of either functional class: *p*0_3_ = 1 – *pSt* – *pSw*.

This functional cell property causes cells to either switch or maintain their preference from their initial preference (during the A/X presentation). In our model, the stable/switching parameter was paramount and so it forced selectivity if a neuron had no response to a stimulus. For example, if a stable cell is selective for A or X, but not B or Y (*p*0_2_ = 1 – *pB* – *pY*), its response will still be selective in both stimulus periods. Note, a cell, that is neither stable or switching, may exhibit responses based only on preference (i.e. *pApBp*0_3_ would contribute to the *pAB* response). Also, a cell functional property can align with its preference (i.e. *pApBpSt* would contribe to the *pAB* response). Thus, under the Stable & Switching model (Fig. S7C), we can write the probability to observing an AB response:

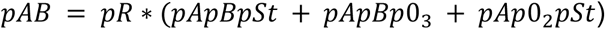

Note, to keep consistency with the independent model, we add a responsiveness parameter (*pR* and *pNR*). A similar equation may be written for the other stable response:

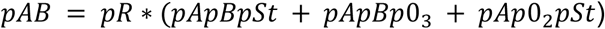

Similar to being stable (*pSt*), cells may exhibit a switching contextual preference (*pSw*). As for stable neurons, when a cell does not have a preference for the second stimuli (*p*0_2_=1 – *pB* -*pY*), the switching property will cause the cell to switch its contextual preference (i.e. AY or XB). Therefore, the probability of observing switching responses can be written as:

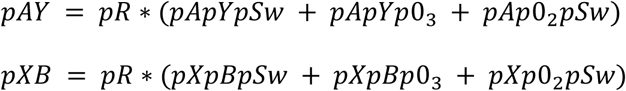

However, it is also possible that the cell’s functional preference does not align with its stimulus selectivity. For example, a neuron might prefer A and B, but have a ‘switching’ functional cell-type. In this case, we assume these properties cancel one another, so no response is observed during the second period (i.e. giving an A0 neuron). Finally, a neuron may have no response during the second period, if it does not prefer B or Y, and also is not a stable or switching neuron. Therefore, the probability of an initial response, followed by no response can be written as:

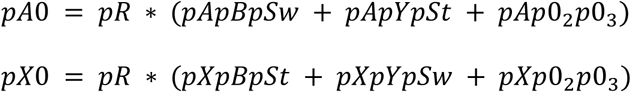

We assume that the functional properties (stable and switching) do not affect response, if the cell is not initially selective (as above, captured by *p*0_1_ = 1 – *pA* – *pX*). Thus, the probability of no initial response to A/X, followed by a response to B/Y is similar to the independent model.

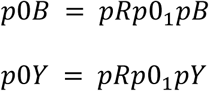

Finally, like in the independent model, observing no response in either stimulus period, can occur, either because the cell is not responsive, or because the cell does not prefer any of the presented stimuli. Again, the functional properties of the cell would not affect the cell’s response. Together, this allows us to write the probability of observing no response, across both the A/X and B/Y periods, as:

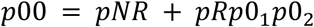

As with the Independent model, we created a simplified version of the stable and switching model, in which the probability of all stimulus preferences was the same (*pA* = *pB* = *pX* = *pY* = *pS*). In this model, we also assumed that a cell is either stable or switching (i.e. it cannot be neither). Thus, we can write *pSw* = 1 –*St*. A summary of both the full ‘Stable & Switching’ model and the ‘Simplified Stable & Switching’ model can found in Fig. S7C and D.

| **Parameter** | **Name** | **Details** |
| --- | --- | --- |
| $pNR$ | Probability no response | |
| $pR$ | Probability response | $pR = 1 - pNR$ |
| $pS$ | Probability of stimuli preference | $pS = pA = pB = pX = pY$ |
| $pA$ | Probability of preferring A | |
| $pX$ | Probability of preferring X | |
| $p0_1$ | Probability of other preference during A/X period | $p0_1 = 1 - pA - pX$ |
| $pB$ | Probability of preferring B | |
| $pY$ | Probability of preferring Y | |
| $p0_2$ | Probability of other preference during B/Y period | $p0_2 = 1 - pB - pY$ |
| $pSt$ | Probability of stable preference | |
| $pSw$ | Probability of switching preference | |
| $p0_3$ | Probability of neither stable or switching selectivity | $p0_3 = 1 - pT - pSw$ |

To compare these two models (and their associated hypotheses), we fit them to the observed response counts across all recorded neurons. We began by determining the selectivity for neurons during the A/X (0-100 ms) and B/Y (175-275 ms) time periods. To do this, we tested if the z-scored firing rate difference was significant (p-value ≤ .025, Bonferroni corrected) during each stimulus presentation period (A/X and B/Y). During the A/X period, cells could be selective to A, X or neither (labeled as 0, when not significant). Likewise, during the B/Y period, cells could be selective to B, Y, or neither (labeled as 0, when not significant). A cell could therefore exhibit one of 9 responses across both time periods (AB, A0, AY, 0B, 00, 0Y, XB, X0, XY; Fig. S8A). For fitting, we divided each count by the total number of counts; this allowed us to fit each probability parameter described above.

Using the minimize function from the scipy.optimize package, we fit each model by minimizing the sum of squared differences across the 9 observed response percentages. For all models, we bound all probabilities to be between (0,1) inclusive. When fitting models, in which all selectivities were equal, we constrained *pS* to (0, 0.5) inclusive (i.e. for the ‘Simplified Independent’ and ‘Simplified Stable & Switching’ models). Additionally, when applicable, we constrained the sum of probabilities, during a given stimulus period, to be less than or equal to 1 (i.e. *pA* + *pX* ≤ 1). Tolerance was set to 1e-100. Other parameters were left to the function’s defaults.

To determine how well each model fit the observed data, we calculated the r^2^ and adjusted r^2^ (which accounts for the number of parameters, (Theil 1961)) for each model:

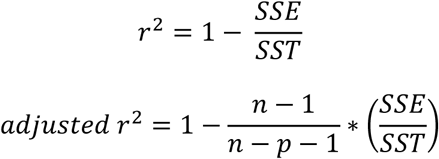

Where, SSE is the sum of squared errors 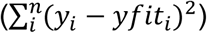, SST is the sum of squared errors across all observations 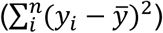 are selectivity probability fits (e.g. *pAB*), *y*_*i*_ are observations,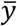 is the mean of observations, *n* is the number of observations, and *p* is the number of parameters. Figure S8B shows both r^2^ and adjusted r^2^.

We found that the Stable & Switching models provided a better fit to the observed cell selectivity profiles (Fig. S8A). The adjusted r^2^ was 0.9988 and 0.9904 for the Stable & Switching and Simplified Stable & Switching models, respectively. This was greater than the Independent and Simplified Independent models (which had adjusted r^2^ of 0.92 and 0.94, respectively). These results suggest the stable and switching dynamics observed are not a result of combining independent selectivities to each chord. Instead, the observed distribution of selectivity across neurons support the hypothesis that there are two classes of neurons: one that stably maintains their contextual preference over the sequence and one that switches their preference during the sequence.

#### The Likelihood of Stable and Switching Neurons were Significantly Greater than Zero

Finally, we wanted to test whether the stable (*pSt*) and switching (*pSw*) parameters were significantly greater than zero. Although the Stable and Switching model fits better than alternatives, this does not necessarily mean that both cell populations exist. To test this in the model, we compared the observed parameter to a null distribution. The null distribution was created by fixing each parameter of interest (*pSt* and *pSw*) to zero. Then, using the Stable & Switching model, we simulated observations for each cell type in the 3×3 table of cell counts (n=999). For each of these “null” cell counts, we refit the full model (no longer holding the *pSt* or *pSw* terms to zero). Thus, the refits of the model create a null distribution of the parameter of interest (1000^th^ value is original parameter fit). Using this distribution, we determined the probability of our observed parameter fit (stars in S8C-F). Importantly, for both the Stable & Switching model and the Simplified Stable & Switching model, the estimated probability of being a stable or switching neuron was significantly greater than zero (p≤1/1000 for both; Fig. S8E and F).

**Figure S1.**
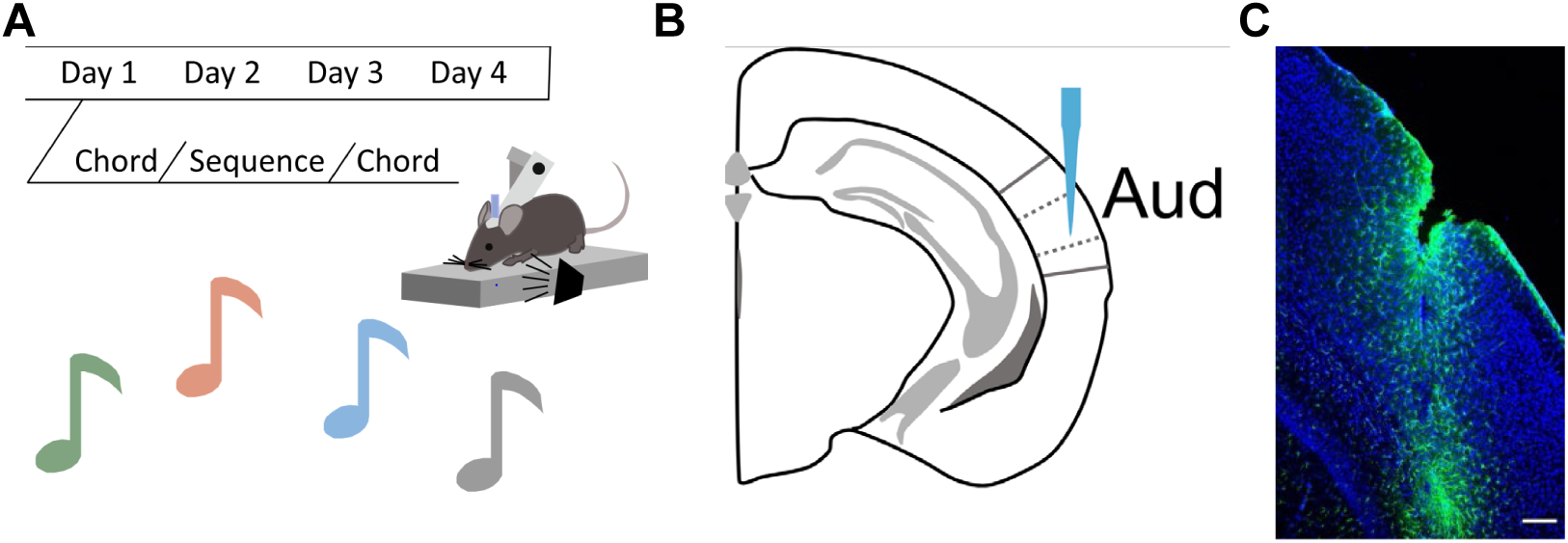
**(A)** Schematic of implicit learning paradigm. On each day, animals heard 300 chords (C/C* in equal proportion), then 1500 sequences (see Fig. 1A for statistics), finishing with 300 chords (C/C*). **(B)** Schematic of electrode location. Silicon probes were implanted in right auditory cortex (stereotaxic coordinates from bregma: −2.7 AP and 4.8 ML). **(C)** Example histology of electrode location. Confocal image taken of cortex around auditory area as shown in B. The scale bar is 150 µm. Green is GFAP (Glial fibrillary acidic protein) immunolabel. Blue is Hoechst stain.

**Figure S2.**
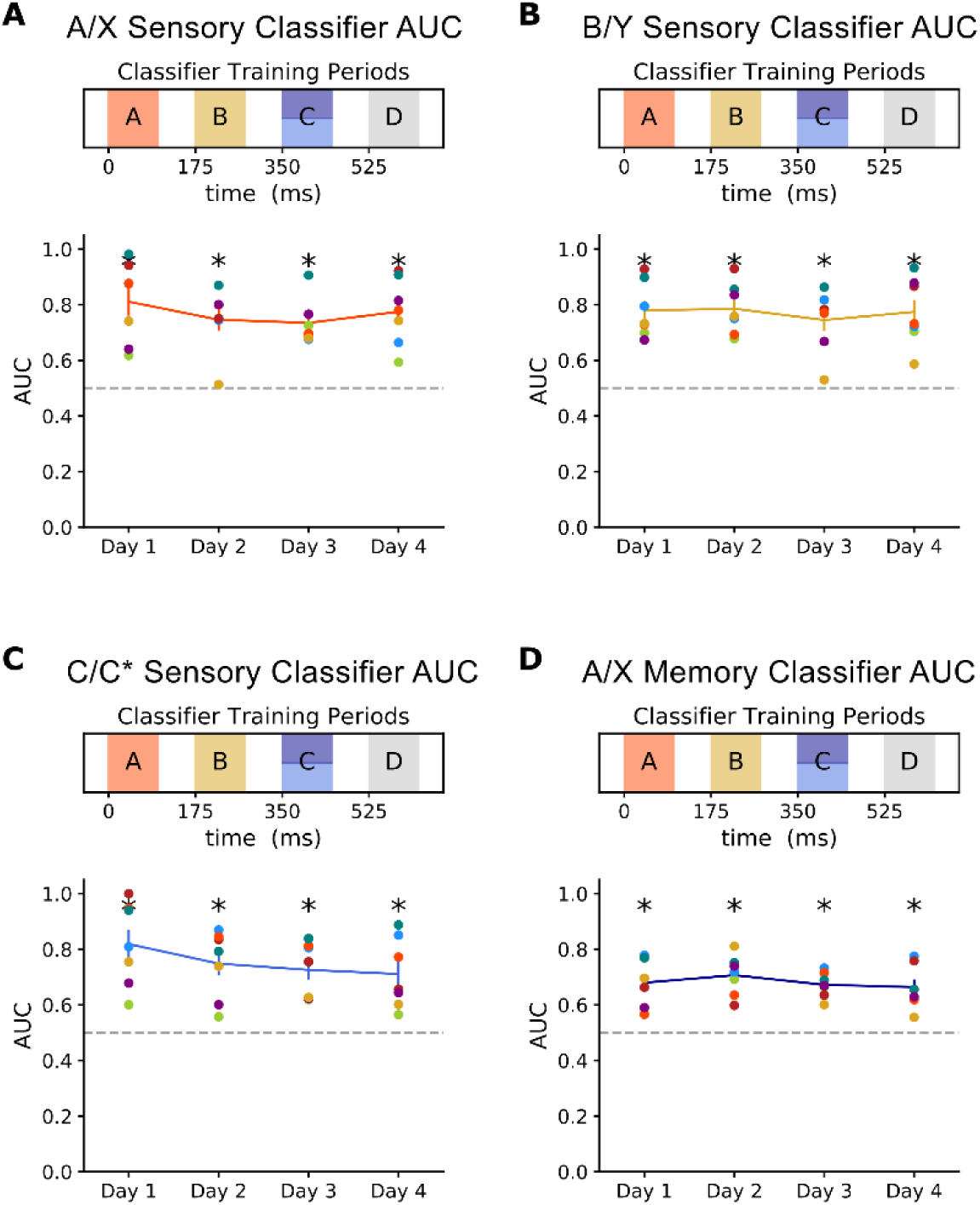
All four classifiers accurately decoded stimulus representations. **(A)** Classifier accuracy for the A/X sensory classifier. Accuracy was measured with AUC (see SI for details). A/X training time was 10-110 ms (orange on timeline). The AUC for each mouse is shown as individual points. Mean and s.e.m across mice is shown as a line across days. Significant differences (p ≤ 0.05) from chance performance (AUC=.5) is indicated with a black star (all were significant). **(B)** Classifier accuracy for the B/Y sensory classifier. B/Y training time was 185-285 ms (yellow on timeline). **(C)** Classifier accuracy for the C/C* sensory classifier C/C* training time was 360-460 ms (light blue on timeline). **(D)** Classifier accuracy for the A/X memory classifier. A/X memory training time was 360-460 ms (dark blue on timeline).

**Figure S3.**
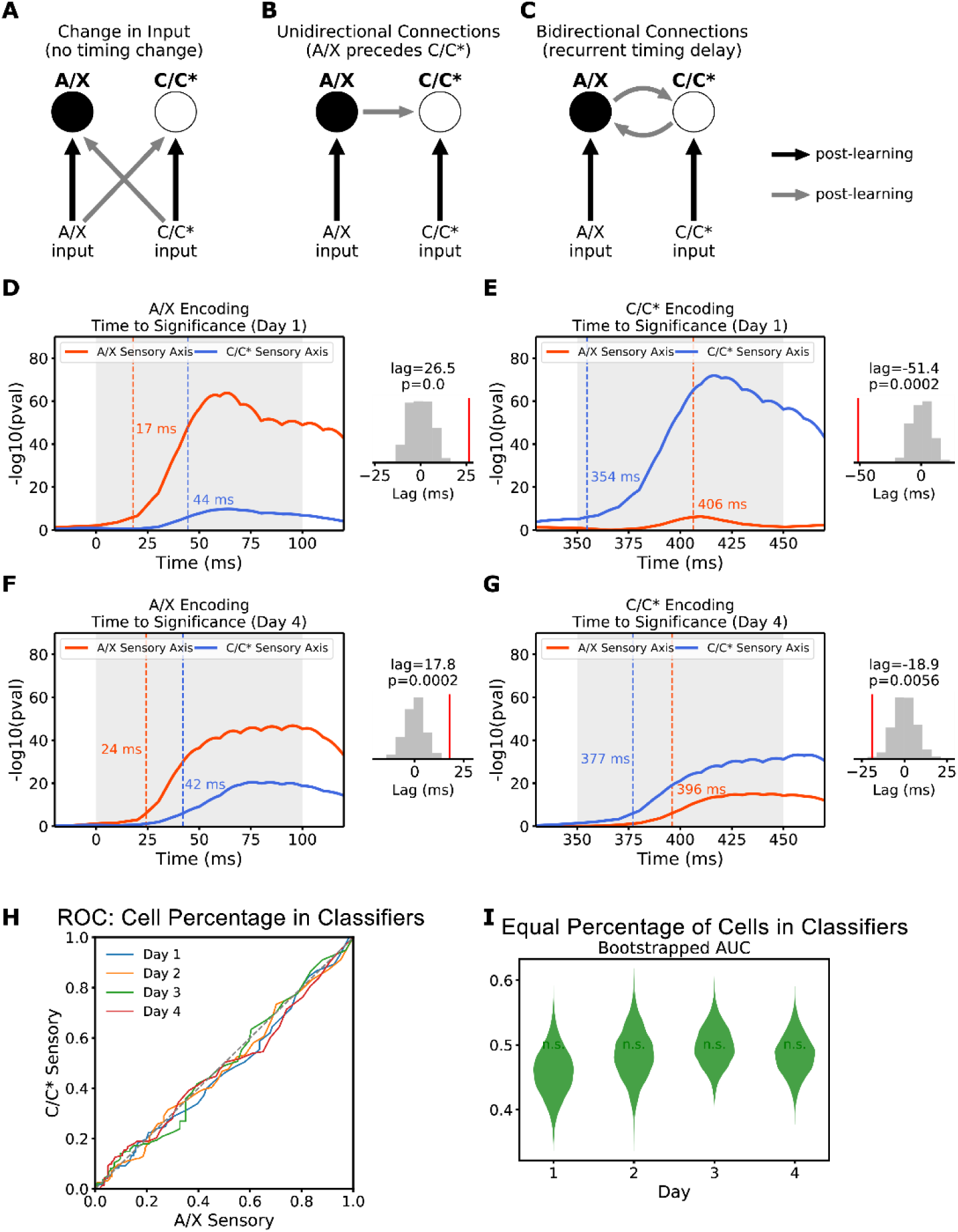
**(A-C)** Schematics of three putative mechanisms for aligning the A/X and C/C* sensory representations. **(A)** The inputs to A/X and C/C* selectivity neurons change, such that associated stimuli induce similar responses within auditory cortex. Such a mechanism does not predict timing differences between A/X and C/C* responses. **(B)** Unidirectional lateral connection from A/X to C/C* representations. This mechanism predicts 1) A/X timing should precede C/C* and 2) a larger percentage of cells should be involved in the A/X representation, because C/C* become active during the A/X presentation. **(C)** Bidirectional lateral connections between A/X and C/C* representations. This mechanism predicts a recurrent timing delay (e.g. A/X precedes C/C* during A/X presentation and vice versa). **(D)** Time to reach significant A/X (orange) and C/C* (blue) encoding during A/X presentation on Day 1. Statistical significance of encoding is on y-axis (−log(p-value)). Vertical dashed lines indicate time to significance along each encoding axis (p ≤ .001, Bonferroni corrected). Inset shows the observed difference in times (the ‘lag’, red vertical line) and their null distribution (grey histogram). **(E)** Time to reach significant A/X (orange) and C/C* (blue) encoding during C/C* presentation. **(F)** Same as **D**, but for Day 4. **(G)** Same as **E**, but for Day 4 **(H)** Receiving operator characteristic (ROC) of percent of cells in each classifier (A/X and C/C* sensory axes). Each day is plotted by a separate line (D1=blue, D2=orange, D3=green, D4=red). **(I)** The percent of cells participating in the A/X and C/C* sensory classifiers on each day were not significantly different. Violin plot shows the distribution of AUC, calculated by bootstrapping the ROC curves.

**Figure S4.**
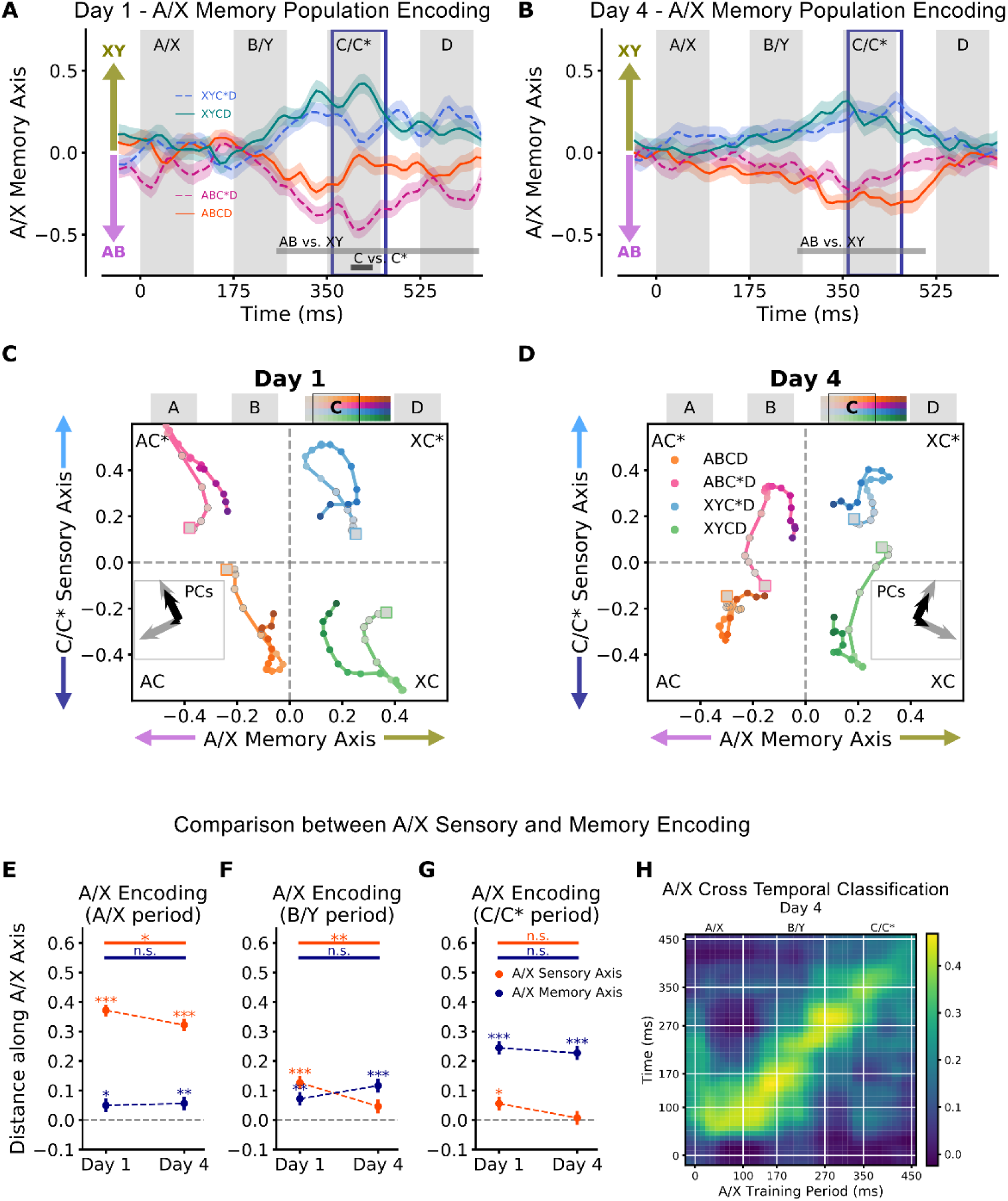
**(A, B)** A/X memory encoding for **(A)** Day 1 and **(B)** Day 4. Firing rate activity was projected onto individual animal A/X memory axes (training period outlined in blue box), z-scored and combined across animals. Average and s.e.m of projections shown across time. Positive projections and negative projections indicate XY (light green) and AB (light purple) encoding, respectively. Light and dark grey bars mark significant differences for AB vs XY and C vs C* respectively (p ≤ .001, Bonferroni corrected t-test). **(C, D)** Neural activity projected into A/X memory - C/C* state space for **(C)** Day 1 and **(D)** Day 4. The x-axis is the projection of neural activity onto the A/X memory axis; the y-axis is the projection onto the C/C* sensory axis. Activity is shown from around the C/C* stimulus period (340 to 520 ms). Marker saturation increases with time. Inset shows PCs of neural trajectories in grey, black arrow size matches percentage of explained variance per PC. **(E-G)** Comparison between A/X sensory (orange) and memory (blue) encoding during the three stimulus periods of the sequence. Mean and s.e.m. of correct distance along each axis combined across trials. To combine across conditions, negatively encoded conditions are flipped. Thus, positive values indicate correct encoding strength. Significant difference between D1 and D4 shown by horizontal bars (permutation test). **(E)** Neural activity from the A/X period (10-110 ms). During the A/X period, A/X sensory encoding is stronger than A/X memory encoding on all days (p≤1/5000, permutation test). **(F)** Neural activity from the B/Y period (180-280 ms). During the B/Y period, on Day 1 A/X sensory encoding is slightly stronger than A/X memory encoding (p=0.042, permutation test), while on subsequent days A/X memory encoding is stronger than A/X sensory encoding (D2 p=0.0002, D3 p=0.022, D4 p=0.021, permutation test). **(G)** Neural activity from the C/C* period (360 – 460 ms). During the C/C* period, the A/X memory encoding was stronger than A/X sensory encoding on all days (p≤1/5000). **(H)** Cross-temporal performance of A/X classifiers. A series of A/X classifiers were trained across the sequence (25 ms windows, shifted by 10 ms steps). Color indicates the encoding strength (withheld data only) for all combinations of training times (x-axis) and test times (y-axis). White bars indicated timing of A/X, B/Y, C/C* periods (and 75 ms inter stimulus intervals). Note the low cross-temporal decoding performance reflects the temporal dynamics of the representation of A/X during the sequence. For all panels, p-values: * ≤ 0.05, ** ≤ 0.01, *** ≤ 0.001

**Figure S5.**
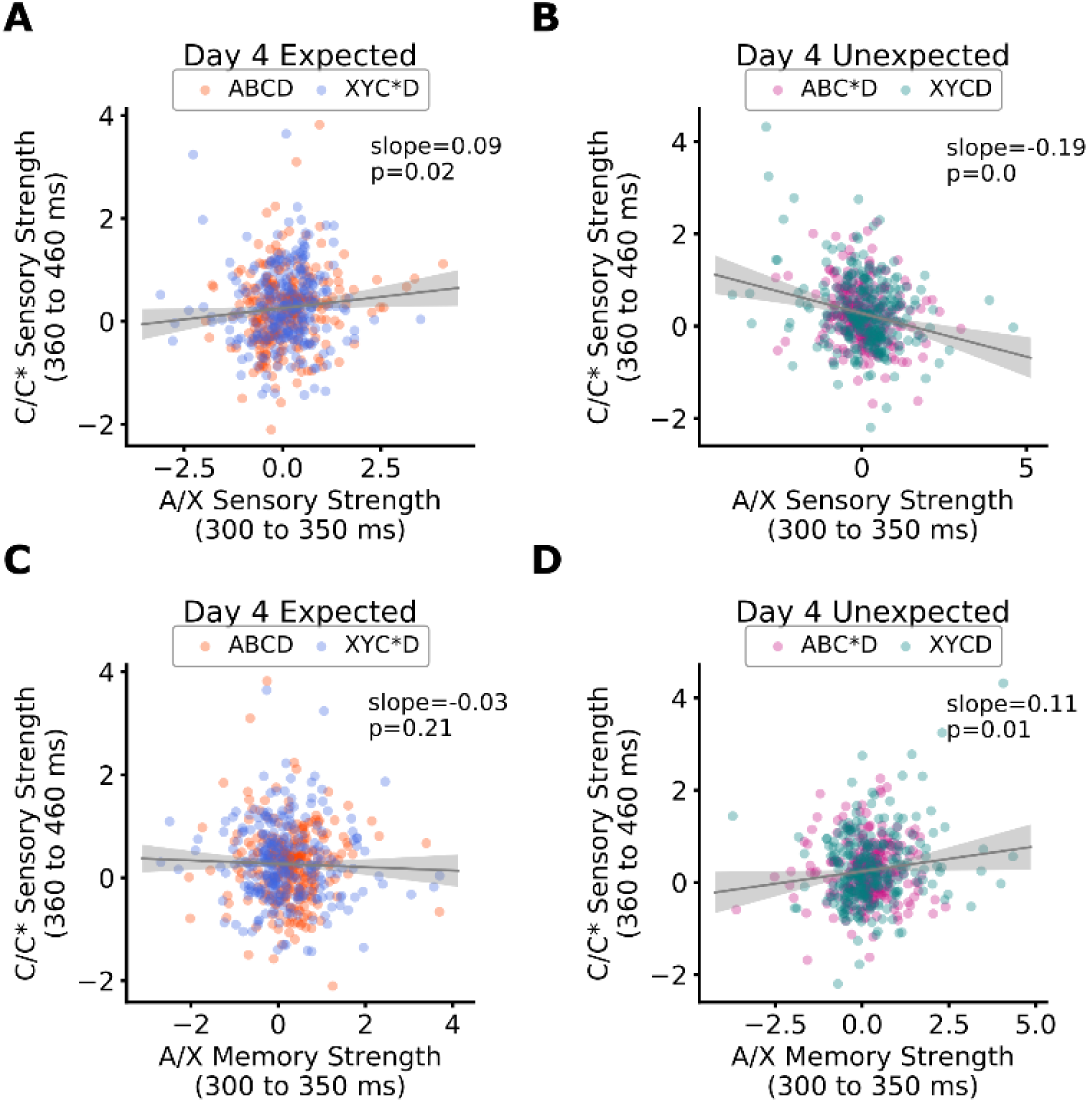
The strength of C/C* sensory encoding is correlated with the strength of both the A/X sensory and memory encoding. All plots show the relationship between C/C* sensory encoding strength (y-axis) and A/X encoding strength (x-axis). Positive and negative values indicate encoding along axis in correct and incorrect directions, respectively. A/X encoding estimated from the 50 ms prior to C/C* onset (300-350 ms). C/C* encoding is estimated from the C/C* period (360-460ms). **(A)** A/X sensory encoding correlates with C/C* encoding accuracy on expected trials (ABCD, XYC*D; bootstrapped linear regression). **(B)** A/X sensory encoding negatively correlates with C/C* encoding accuracy on unexpected trials (ABC*D, XYCD). **(C)** A/X memory encoding accuracy is weakly negatively correlated with C/C* encoding on expected trials. **(D)** A/X memory encoding accuracy is positively correlated with C/C* encoding during unexpected trials.

**Figure S6.**
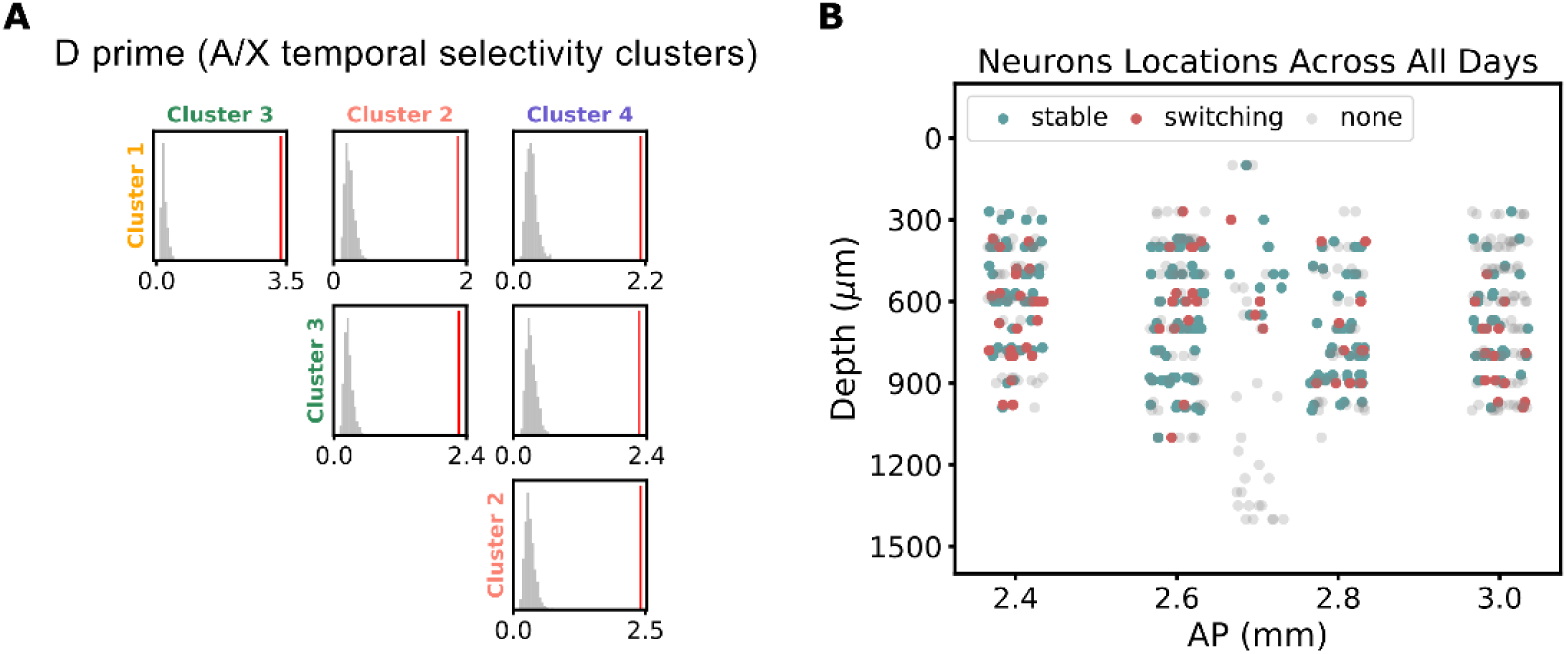
**(A)** Sensitivity index (d-prime) calculated between clusters. Four clusters of neuron temporal selectivity were found using a Phenograph algorithm (Fig. 4, see SI for details). The d-prime between all pairs of cluster groups was calculated (red line); grey distributions are null d-prime distributions, estimated with a permutation test (1000 shuffles). Stable cells were clusters 1 and 3, switching cells were clusters 2 and 4. **(B)** Estimated locations of functional clusters along recording arrays. Switching (red), stable (green), and none (grey) cells are plotted according to their estimated electrode location (x-axis – AP, y-axis – depth (DV) based on implant coordinates; 6 probes had 4 shanks separated by 200 µm). Small, random jitter in anterior-posterior (AP) direction is added for clarity of presentation and do not reflect actual differences.

**Figure S7.**
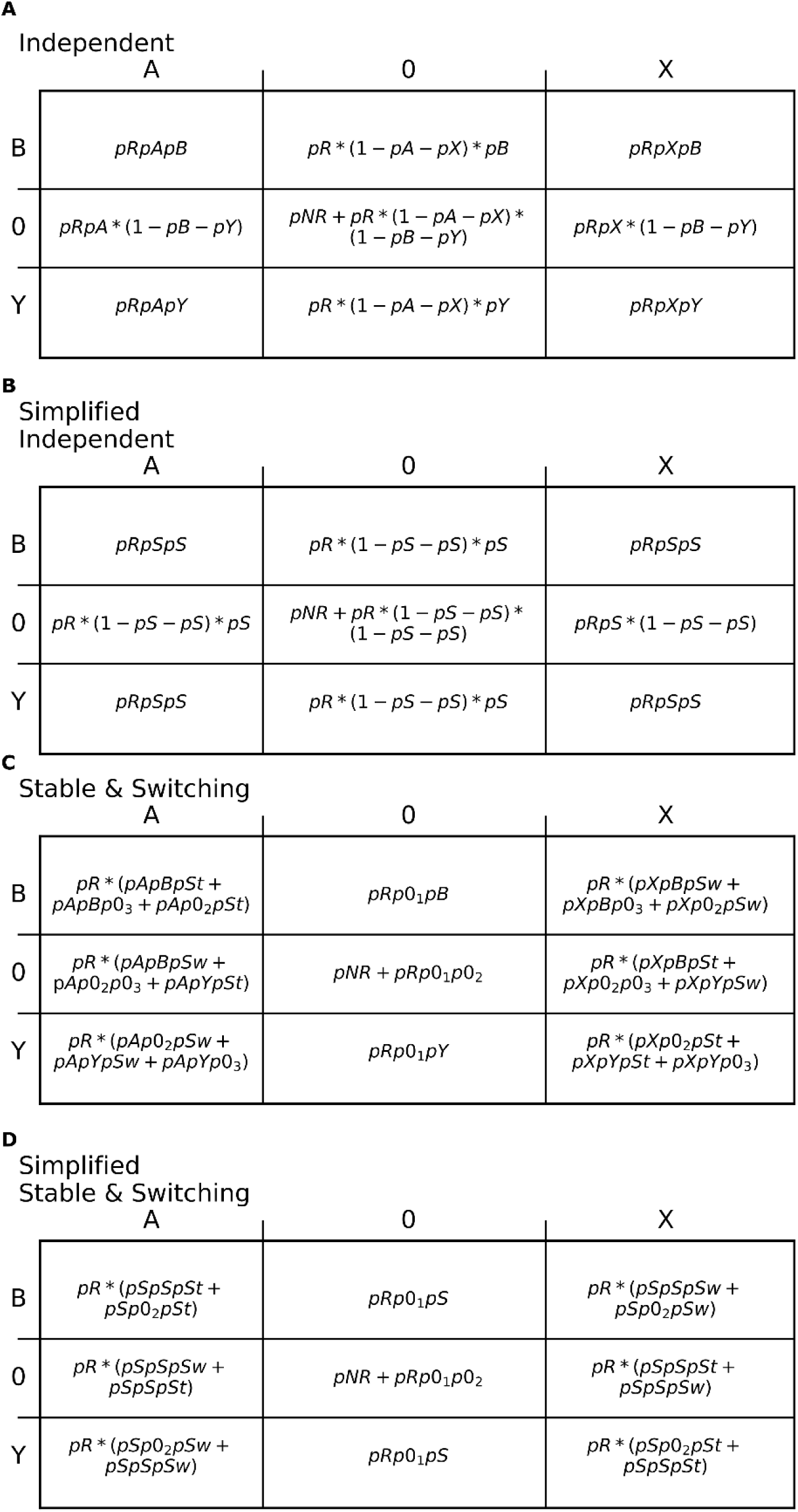
Table of probabilities for each stimulus response pair across A/X and B/Y periods for all models. **(A)** Independent model. **(B)** Simplified Independent model. **(C)** Stable & Switching model. **(D)** Simplified Stable & Switching model.

**Figure S8.**
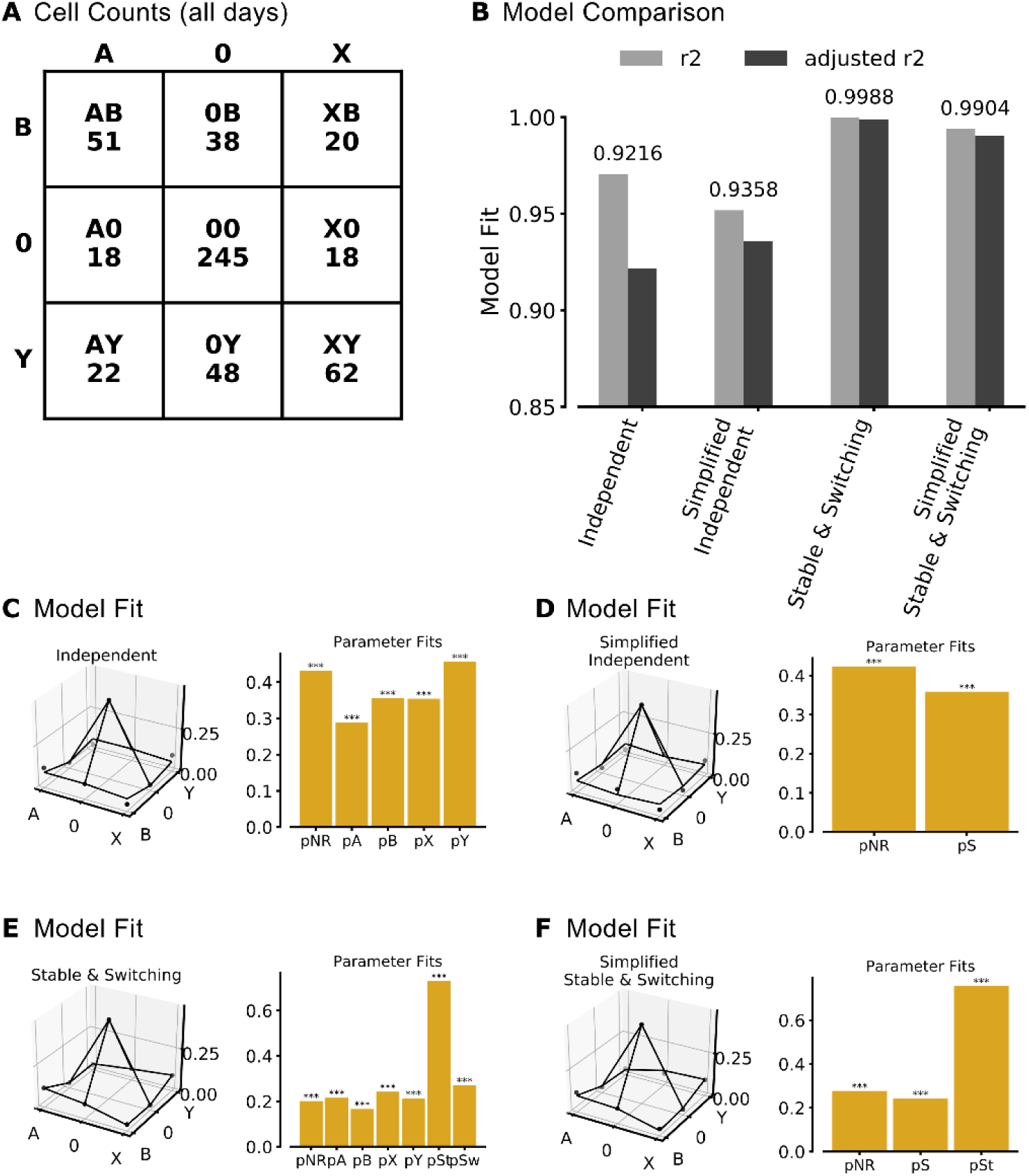
**(A)** Counts of cells with a specific stimulus preference across the A/X and B/Y stimulus periods. Columns indicate A/X period preference (A, 0 (none), or X). Rows indicate B/Y period preference (B, 0 (none), or Y). Each box contains selectivity label (e.g. AB) and corresponding observed cell count (across all days). **(B)** Model comparison across Independent and Stable & Switching models. R^2^ (grey) and adjusted r^2^ (black) calculated per model. The number above each set of bars / model is adjusted r^2^. Both Stable & Switching models have higher adjusted r^2^ (0.99, 0.99) compared to the Independent models (0.92, 0.94). **(C, D)** Parameter fits for the **(C)** Independent model and **(D)** Simplified Independent model. Left panel: observed selectivity probabilities (dots) and model fit (lines). X-axis is selectivity during A/X period; y-axis is selectivity during B/Y period; z-axis is percent of cells with each combined selectivity type. Right panel: parameter fits for each probability. Stars indicate the probability of the observed value being greater than zero. **(E,F)** Same as (C,D) but for the **(E)** Stable & Switching model and the **(F)** Simplified Stable & Switching model.

